# Moral Hazard Heterogeneity: Genes and Health Insurance Influence Smoking after a Health Shock

**DOI:** 10.1101/2021.03.05.434163

**Authors:** Pietro Biroli, Laura Zwyssig

## Abstract

Decision-making in the realm of health behaviors, such as smoking or drinking, is influenced both by biological factors, such as genetic predispositions, as well as environmental factors, such as financial liquidity and health insurance status. We show how the choice of smoking after a cardio-vascular health shock is jointly determined by the interaction between these biological and environmental constraints. Individuals who suffer a health shock when uninsured are 25.6 percentage points more likely to reduce smoking, but this is true only for those who have a low index of genetic predisposition to smoking. Individuals with a low index of genetic predisposition are more strategic and flexible in their behavioral response to an external shock. This differential elasticity of response depending on your genetic variants is evidence of individual-level heterogeneity in moral hazard. These results suggest that genetic heterogeneity is a factor that should be considered when evaluating the importance and fairness of health insurance policies.

**JEL CODES:** I12, I13, D63, D91

## 1 Introduction

Moral hazard shapes behaviors in all domains characterized by asymmetric information (Arrow, 1963; Finkelstein, 2014). Propensity to moral hazard can differ across individuals, the same as preferences and beliefs. Unlike preferences and beliefs, heterogeneity in moral hazard has been proposed but never directly measured (Dubois and Vukina, 2009; Einav et al., 2013; Kowalski, 2018). Leveraging information on genetic variants associated with smoking behavior, we provide a novel approach to measuring heterogeneity in moral hazard.

To measure heterogeneity in moral hazard, we estimate how individuals carrying different genetic variants differentially change their smoking behavior following a cardio-vascular health shock when they are not covered by health insurance. Combining the genetic and moral hazard factors shaping health behaviors, we contribute to the burgeoning literature studying gene-environment interactions (Barcellos, Carvalho and Turley, 2018; Belsky et al., 2018; Caspi et al., 2002; Fletcher, 2012; Papageorge and Thom, 2020; Schmitz and Conley, 2017*a*,*b*; Wedow et al., 2018). Specifically, this study is the first to show how genetic risk and insurance coverage jointly influence individual choices of health behavior. We show that moral hazard stemming from health insurance (financial risk) interacts with the genetic predisposition for smoking (Sgenetic risk) in determining the probability of smoking cessation following a health shock. Using data from the Health and Retirement Study (HRS), a longitudinal population-based study of elderly Americans, we estimated how experiencing a health shock between survey waves affected the smoking probability of individuals with different coverage levels and genetic predispositions for smoking.

To cleanly identify this interplay between genes and the environment, we leverage a key feature of the US health insurance system: Medicare, which provides public health insurance to all US citizens older than 65. Before the introduction of the Affordable Care Act (Obama, 2016), a significant fraction of the population younger than 65 was still uninsured (Barnett and Vornovitsky, 2016; Cohen et al., 2009). Exploiting the differential timing of health shocks before or after the Medicare-eligibility age of 65 for previously uninsured individuals, we estimate that Medicare eligibility reduced smoking cessation rates after a health shock by 25.6 percentage points, but only for those individuals with a low index of genetic propensity to smoking. Comparing this effect between individuals with a high versus a low index of genetic predisposition for smoking allows for an assessment of how the moral hazard problem found in previous research interacts with genetic risk.

We focus on smoking behavior because chronic diseases and health care costs caused by tobacco use are estimated to be one of the biggest health challenges in industrialized countries, and have rapidly increased in importance in the developing world (Goodchild, Nargis and Tursan d’Espaignet, 2018; United States Department of Health and Human Services, 2014). In the US, smoking is estimated to cause more than 400,000 premature deaths annually (Ma et al., 2018), and the economic costs of smoking-related illness amount to around $300 billion each year, including almost $170 billion for direct medical care and an additional $156 billion in lost productivity (United States Department of Health and Human Services, 2014; Xu et al., 2015). Both the health burden from smoking-related illness as well as the economic burden on an already strained health care system have made it a priority to understand what factors affect individuals’ smoking decisions, and how health care can effectively encourage cessation.

Smoking cessation, like many other health behaviors, is influenced both by environmental and genetic factors. One environmental factor that has been associated with a reduction in tobacco consumption is health insurance coverage (Dave and Kaestner, 2009; Marti and Richards, 2017; Richards and Marti, 2014), especially after experiencing a severe smoking-related health shock, like the onset of a cardiovascular illness (Clark and Etilé, 2002; Falba, 2005; Keenan, 2009; Khwaja, Sloan and Chung, 2006*a*; Sundmacher, 2012). Alleviating the financial burden of health care costs, health insurance can have the unintended side effect of preventing beneficial behavior changes that would have taken place if the individuals were fully responsible for the financial consequences of poor health (Marti and Richards, 2017). This adverse incentive created by health insurance is a typical example of “moral hazard”: the notion that individuals change their behavior in an undesired way because the consequences of their actions are not (fully) borne by themselves (Einav and Finkelstein, 2018; Zweifel and Manning, 2000).

Another factor that is tightly linked to smoking behavior is genetic makeup. Genetic factors can explain around 30% to 85% of the variance in regular smoking, according to several studies comparing identical and fraternal twins (Boardman, Blalock and Pampel, 2010; Hall, Madden and Lynskey, 2002; Heath et al., 1993; Li et al., 2003; Sullivan and Kendler, 1999). In recent years, significant progress has been made in identifying genetic variants associated with susceptibility to smoking (Liu et al., 2010, 2019; The Tobacco and Genetics Consortium et al., 2010; Thorgeirsson et al., 2010, 2008). In particular, most genetic variants from the *CHRNA5-CHRNA3-CHRNB4* gene cluster, which influences nicotine response and metabolism and which studies link to nicotine dependence, are strongly associated with smoking phenotypes (Stoker and Markou, 2013). Genetic variants near dopamine receptors are strongly associated with daily smoking and difficulty in cessation, but not with smoking initiation-related phenotypes, suggesting that dopamine-related variants become more relevant as an individual’s nicotine use progresses (Liu et al., 2019). Indices of genetic predisposition to smoke have been shown to be related to smoking initiation, and individuals of higher genetic risk are more likely to develop dependence faster and more frequently and to fail in their cessation attempts (Belsky et al., 2013). Thus, genetic data may capture an individual’s predisposition for smoking behaviors via multiple biological channels including addiction in addition to nicotine response.

These developments in mapping the genetic architecture of health behaviors, together with an increased availability of genetic measures in large representative surveys, allow for a better understanding of how genetics can interact with other environmental factors in determining smoking behavior and nicotine dependence. For example, adolescent environmental shocks have been shown to alter the influence of specific genetic variants on an individual’s risk of developing nicotine dependence (Bierut, Johnson and Saccone, 2014; Chen et al., 2009; Johnson et al., 2010). More generally, the relationship between genetic variants and smoking has been shown to depend on neighborhood characteristics (Meyers et al., 2013) the cohort of birth (Domingue et al., 2016; Wedow et al., 2018), military service in the Vietnam era (Schmitz and Conley, 2016), and tobacco taxes (Fletcher, 2012; Slob and Rietveld, 2020).

Our results highlight the importance of considering genetic predisposition when evaluating behavioral responses to shocks and policies, such as health insurance coverage (Harper, 1993; Morrison, 2005). Genetic predispositions can curb the negative behavioral consequences and the moral hazard associated with changes in health insurance status. Besides being relevant for the debate about equal and fair access to health insurance, our results highlight a new avenue of potential future research: leveraging recent advances in molecular and human genetics, we identify a new form of individual heterogeneity in the response to treatments. This heterogeneity used to be unobserved, which could lead to incorrect policy conclusions. In the era of genomics and personalized medicine, an individual genetic makeup can be a factor not only in the treatment they receive, but also in their response to insurance coverage (Khera et al., 2018; Ritz et al., 2017; Schork, Schork and Schork, 2018; Torkamani, Wineinger and Topol, 2018). A new form of individual heterogeneity in treatment effects (Papageorge and Thom, 2020). Since individuals are endowed at conception with their genetic makeup, and they have not done anything to deserve it or be held accountable for it (Barth, Papageorge and Thom, 2019; Harden, In Press; Kweon et al., 2020; Pereira, 2021), this differential response to insurance shocks raises questions of fairness and equality in the public debate over health insurance policies.

More generally, we show how leveraging recent developments in behavioral and molecular genetics can shed new light on fundamental economic concepts, such as elasticity of response to health shocks and heterogeneity in moral hazard. These concepts are formalized more precisely in the model in Appendix E. Indices of genetic predisposition to health behaviors provide readily available individual measures of heterogeneity that can enrich our economic models.

## 2 Data

### 2.1 Study Sample

The HRS is a nationally representative longitudinal household survey initiated in 1992 among US citizens aged 50 and older, followed for 12 biannual waves over 22 years, containing detailed medical, economic, social, psychological, and genetic information about the respondents (Sonnega et al., 2014).^1^

Genetic information comes from DNA samples collected in face-to-face interviews, for which a random subset of HRS households were selected to participate in 2006, 2008, and 2010. Saliva samples are collected and genotyped for more than 15,000 participants (Ware, Schmitz and Faul, 2017). More information on the DNA extraction and the genotyping process is provided in Section C.1 in the Appendix.

The analysis is done on a subsample of the data selected based on criteria of age (between 60-70 at the time of interview), ever-smokers at baseline (their first observation), observed in at least 2 different time periods, and genetically of European descent. 5,854 HRS respondents satisfy these criteria and have non-missing values for the main variables of interest (smoking status, polygenic score (PGS) for regular smoking, insurance status, and the occurrence of heart conditions or strokes).^2^ The age restriction imposed on the study sample increases comparability between those experiencing a health shock before or after age 65. The restriction imposed on genetic ancestry is best-practice in social-science genetics to avoid issues of population stratification, for example, mistaking a genetic variant with different allele frequencies in Asians and Caucasians for a marker of ability to use chopsticks (Hamer and Sirota, 2000). The restriction also increases the sample’s similarity with the genetic profile of the discovery sample used in the genome-wide association study (GWAS) that informed the weights for the Single Nucleotide Polymorphisms (SNPs) used for calculating the PGS (Freese, 2018; Martin et al., 2017).^3^ Also, we exclude respondents who reported the onset of a cardiovascular illness since the last survey wave when interviewed at ages 65 or 66. Since HRS surveys are only conducted every 2 years, it is not possible to determine Medicare eligibility status at the time of the health shock for these individuals.^4^

### 2.2 Variables of interest

#### Genetic propensity

As a measure of an individual’s genetic propensity to smoke, we leverage recent developments in molecular and behavioral genetics to construct an index known as a polygenic score (PGS). A polygenic score is a weighted average of the number of risky genetic variants *G_ij_* associated with the probability of smoking that are carried by a particular individual. For our measure of genetic variants *G_ij_*, we follow the literature and use the most common form of genetic variation: Single Nucleotide Polymorphisms (SNP), a one base-pair substitution at a particular location (locus) on the human genome.

Specifically, the scores are calculated as follows:

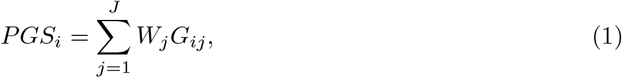

where *G_ij_* is the genotype for individual *i* at each of the more than 10 million SNP *j*; the weight *W_j_* is the effect size for SNP *j* estimated via meta-analysis in a different sample than the HRS by the GWAS and Sequencing Consortium of Alcohol and Nicotine use (GSCAN) (Liu et al., 2019). The scores have been normalized to have a mean of zero and a standard deviation of one.^5^

To avoid functional form assumptions and maximize power in the statistical analysis, the sample is divided into high- and low-genetic propensity to smoking based on the PGS. A high-PGS indicator is defined to take the value 1 for individuals in the top 2/3 of the PGS distribution.^6^ As shown in Figure 1, the distribution of the PGS is bell-shaped, it is shifted to the right for baseline smokers, and it is predictive of current smoking behavior: ≈ 30% of individuals with a PGS higher than 2.5 are current smokers, while only ≈ 10% of individuals with a PGS lower than −2.5 currently smoke.

**Figure 1:**
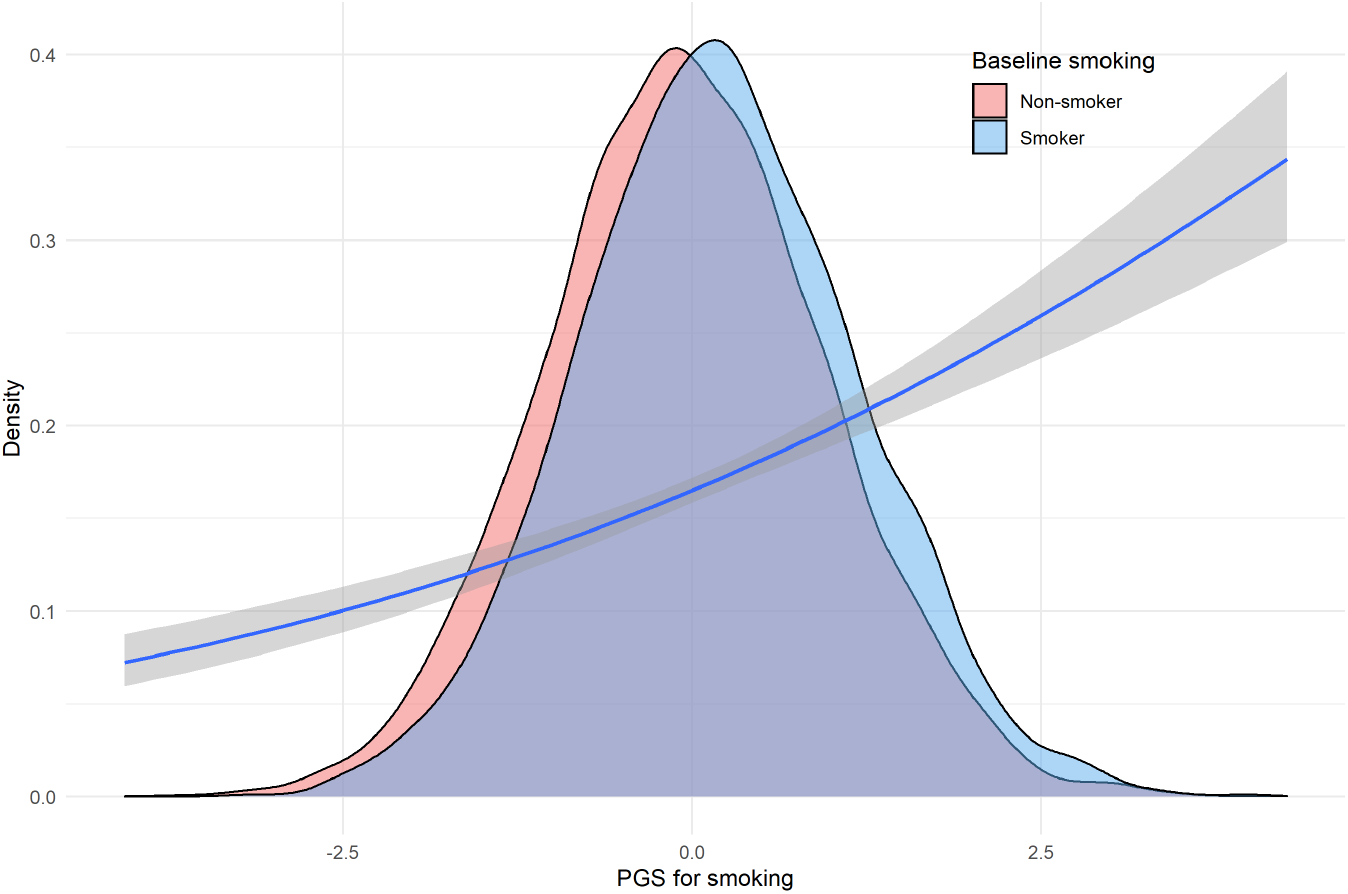
PGS distribution and correlation with smoking behavior. *Notes:* Distribution of Polygenic Score (PGS) for baseline smokers (blue) and non-smokers (red). Generalized linear smoothed correlation between current smoking and PGS shown in the blue line (with 95% confidence intervals in shaded grey area). *Data used*: HRS waves 1-13, restricted to observations with age between 60 and 70 years.

#### Health shocks

Following previous studies (Falba, 2005; Keenan, 2009; Khwaja, Sloan and Chung, 2006*a*,*b*; Smith et al., 2001), we focus on the first o ccurrence of an acute cardiovascular condition. Cardiovascular conditions include either a heart problem (heart attack, coronary heart disease, angina, congestive heart failure, or other heart problems) or a stroke (or transient ischemic attack). These conditions have strong links to tobacco use, have a relatively high incidence among older adults, and are likely to require immediate and intensive use of costly medical services (Lloyd-Jones et al., 2010; Teo et al., 2006; Thorpe, Florence and Joski, 2004). Additionally, they can occur repeatedly for the same individual and thereby incentivize improvements in health behaviors to prevent a recurrence.

The health shock is a binary indicator set to 1 if the respondent is diagnosed with a new cardiovascular condition during the time since the last HRS survey, but does not have a history of cardiovascular disease prior to this diagnosis. The exact timing of the health shock within the past 2 years since the last interview is not observed, and the diagnoses are all self-reported.

As shown in Figure 2, the rate of cardiovascular health problems increase with age from about 2% to 4% annually, but it is not strongly related to the PGS and, most importantly, does not jump around the age of 65 (see also Figure 11 in the Appendix).

**Figure 2:**
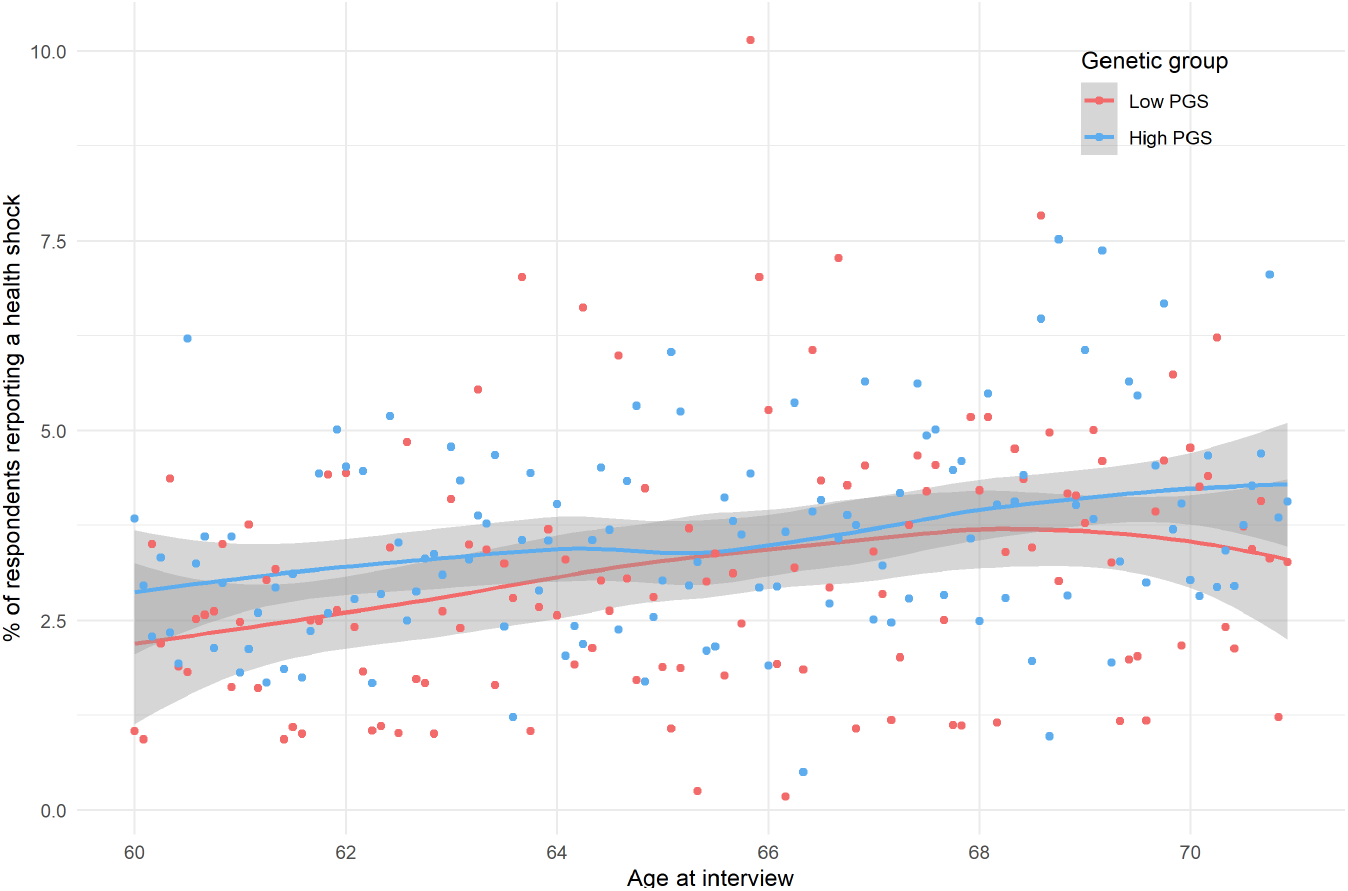
Prevalence of cardiovascular health shock. *Notes:* Self-reported indicator of having been diagnosed for the first time with a cardiovascular condition since the last HRS survey. Age refers to the time of the survey, not the time of the health shock, which is unknown up to a 2-year windows, since HRS surveys are bi-annual. Bin-scattered plot and generalized linear smoothed correlation between age at interview and cardiovascular health shock shown in red (low PGS = lowest tertile of polygenic score) and blue (high PGS = upper two tertiles of polygenic score). *Data used*: HRS waves 1-13, restricted to observations with age between 60 and 70 years.

#### Health Insurance

Before the age of 65, HRS respondents are classified as uninsured in a given survey wave if they report not being covered by *any* health insurance plan. Respondents are classified as being *persistently uninsured* if they were uninsured in every wave between 60 and 65.

The share of persistently uninsured individuals is shown in panel A of Figure 3.

**Figure 3:**
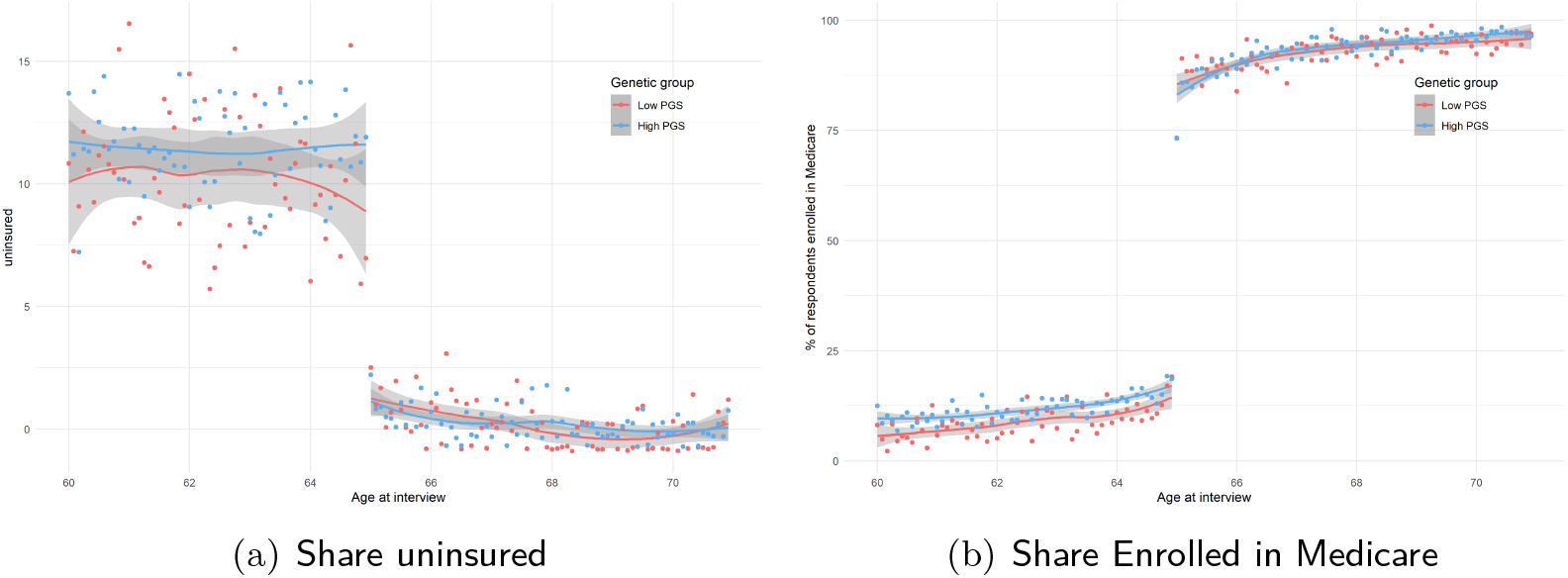
Share uninsured and enrolled in Medicare across the age 65 threshold by PGS. *Notes:* Self-reported indicator for lack of any insurance coverage and being enrolled in Medicare. Bin-scattered plot and generalized linear smoothed correlation between age and outcome variables shown in red (low PGS = lowest tertile of polygenic score) and blue (high PGS = upper two tertiles of polygenic score). *Data used*: HRS waves 1-13, restricted to observations with age between 60 and 70 years.

After the age of 65, individuals are considered eligible for Medicare. As shown in Panel B of Figure 3, the vast majority of individuals are aware of Medicare eligibility: more than 90% of individuals report being covered by Medicare after the age of 65.

#### Smoking status

The outcome of interest is a self-reported binary indicator for current smoking status. It is set to 1 if the respondent is an ever-smoker (having smoked more than 100 cigarettes throughout one’s life) and smoking at the time of the survey.

As shown in Figure 4, the prevalence of smokers in the sample decreases with age and is about 5 percentage points higher for the individuals with a high PGS.

**Figure 4:**
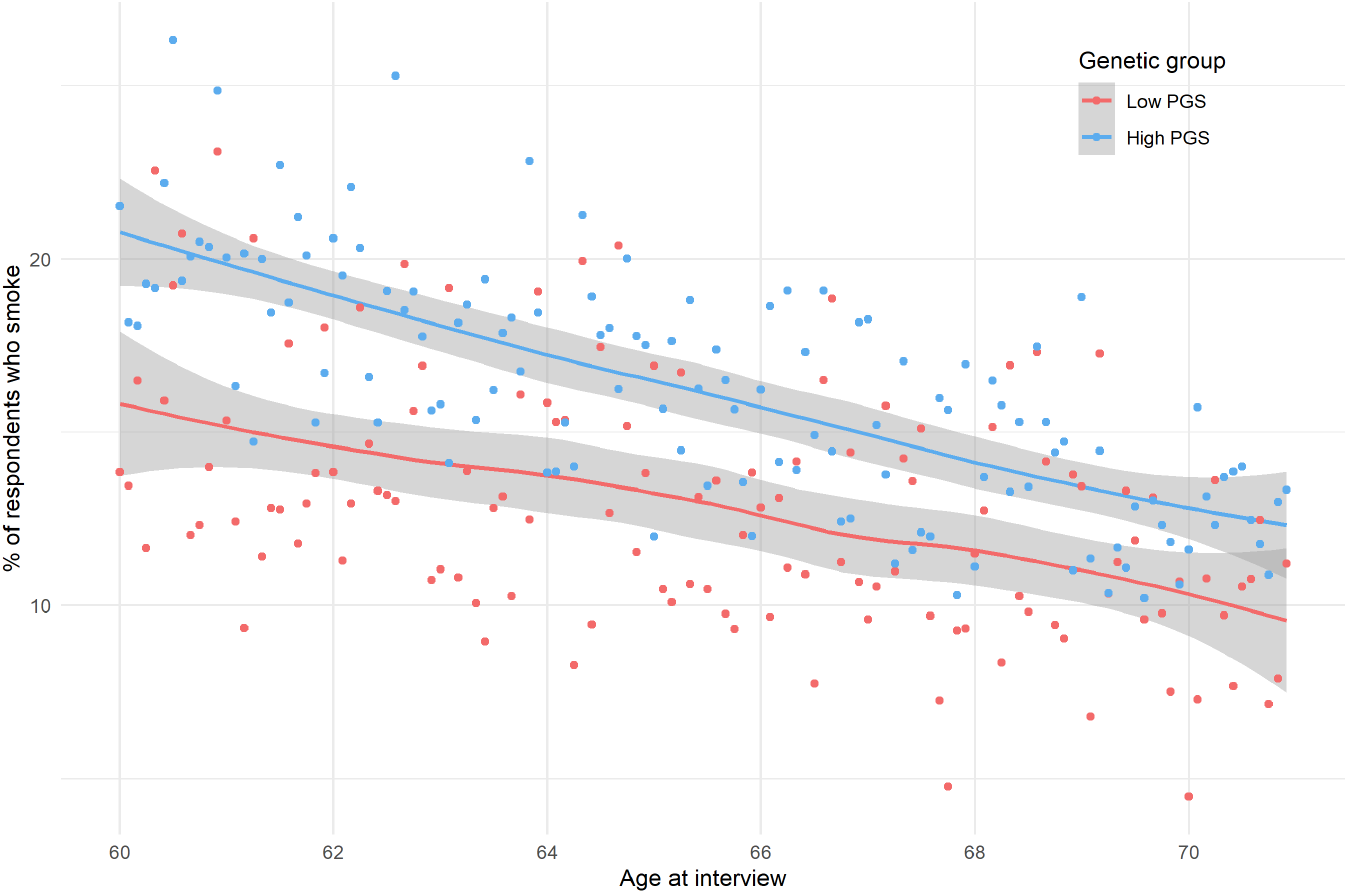
Prevalence of smoking. *Notes:* Self-reported smoking rate by age and high or low PGS. Bin-scattered plot and generalized linear smoothed correlation between age and smoking shown in red (low PGS = lowest tertile of polygenic score) and blue (high PGS = upper two tertiles of polygenic score). *Data used*: HRS waves 1-13, restricted to observations with age between 60 and 70 years.

More information on the construction of all variables used in the analysis can be found in Section C.3 in the Appendix.

### 2.3 Sample Characteristics

Characteristics of the study sample used in the analysis are summarized in the first column of Table 1. The study sample consists of 5,854 individuals (and 26,022 personyear observations). The average age at baseline is 61.2 years, 49.9% of the sample is made up of women, and 29.5% smoke at baseline. Individuals are observed for 4.5 waves on average, a good panel dimension even if restricted to the 60-70 age range. 5.9% of individuals in the sample are uninsured in all observations before the age of 65, and 12.3% experience a cardiovascular health shock during the observation period. The second and third columns in Table 1 describe the sample stratified into 2 genetic groups. High-PGS respondents are 4 percentage points more likely to smoke at baseline, 0.7 percentage points more likely to be uninsured before the age of 65, and included relatively more women compared to the low-PGS respondents.^7^

**Table 1:**
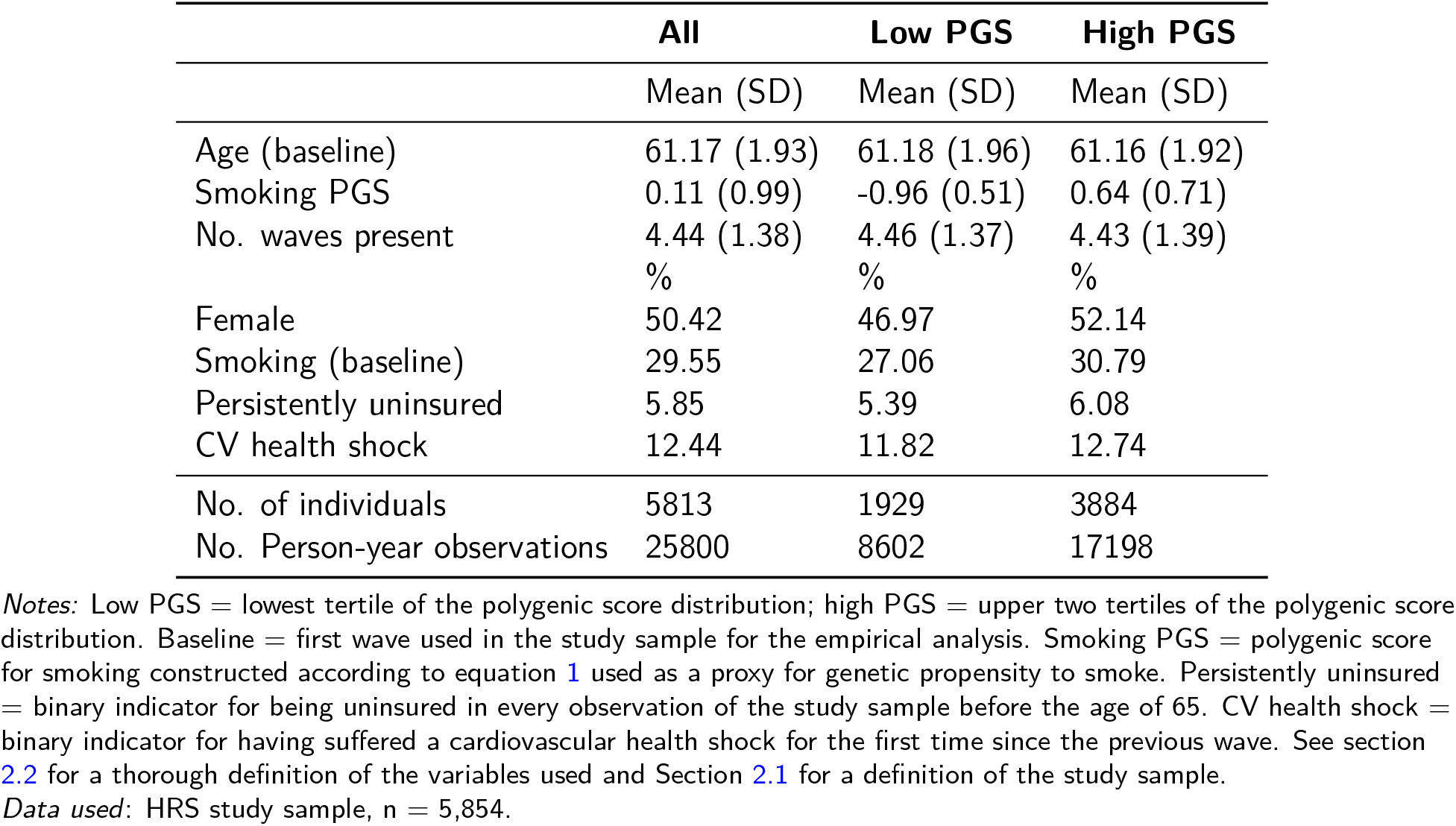
Descriptive Statistics for the Study Sample, Stratified by Genetic Group

|  | All | Low PGS | High PGS |
| --- | --- | --- | --- |
|  | Mean (SD) | Mean (SD) | Mean (SD) |
| Age (baseline) | 61.17 (1.93) | 61.18 (1.96) | 61.16 (1.92) |
| Smoking PGS | 0.11 (0.99) | -0.96 (0.51) | 0.64 (0.71) |
| No. waves present | 4.44 (1.38) | 4.46 (1.37) | 4.43 (1.39) |
|  | % | % | % |
| Female | 50.42 | 46.97 | 52.14 |
| Smoking (baseline) | 29.55 | 27.06 | 30.79 |
| Persistently uninsured | 5.85 | 5.39 | 6.08 |
| CV health shock | 12.44 | 11.82 | 12.74 |
| No. of individuals | 5813 | 1929 | 3884 |
| No. Person-year observations | 25800 | 8602 | 17198 |

Notes: Low PGS = lowest tertile of the polygenic score distribution; high PGS = upper two tertiles of the polygenic score distribution. Baseline = first wave used in the study sample for the empirical analysis. Smoking PGS = polygenic score for smoking constructed according to equation 1 used as a proxy for genetic propensity to smoke. Persistently uninsured = binary indicator for being uninsured in every observation of the study sample before the age of 65. CV health shock = binary indicator for having suffered a cardiovascular health shock for the first time since the previous wave. See section 2.2 for a thorough definition of the variables used and Section 2.1 for a definition of the study sample.
Data used: HRS study sample, $n = 5,854$ .

## 3 Empirical Analysis

In a within-person analysis with individual fixed effects, we compare the change in smoking rates after suffering a health shock across four types of persistently uninsured people: younger or older than 65 (as a proxy of exposure to health insurance via Medicare) and high or low genetic propensity to smoking. We focus on persistently uninsured people, i.e. people who reported being uninsured in every observation of the study sample before age 65, to zoom in on the potential for moral hazard.

The identification strategy builds on previous work (Marti and Richards, 2017) and leverages the differential timing of health shocks before or after Medicare eligibility at age 65. Conditional on having suffered a health shock, the exact *timing* of the shock is arguably both exogenous and unanticipated. While the probability of experiencing a health shock increases with age (Lloyd-Jones et al., 2010), no observable jump or change in trend can be seen in our data in the percentage of respondents reporting a health shock around the age of 65 (see Figures 2 and 11 as well as Section C.4.1 in the appendix for details). Notice, however, that only the age at the time of the interview is known, not the age at the time of the cardiovascular health shock, which could have happened any time since the prior interview (about 2 years before). Therefore, our dataset is not precise enough to estimate a possible discontinuity in the probability of suffering a cardiovascular health shock at age 65. Nevertheless, the absence of a jump at the time of Medicare eligibility is consistent with the work of Card, Dobkin and Maestas (2009, 2008) who, using much more granular data, find a sharp increase in health care utilization at age 65, but no discontinuity in the probability of suffering from a health shock.

The absence of a jump is reassuring for our identification strategy, which relies on a comparison before and after the age of 65: accounting for age, respondents experiencing the shock after 65 should not be systematically different from respondents experiencing the shock before 65. This setting thus provides a good framework for estimating the causal effect of health care eligibility on the smoking response to a health shock in individuals who are uninsured before the age of 65.

Conceptually, this design aims to mimic the following hypothetical comparison: within both the low-PGS and the high-PGS group, compare the change in smoking status after suffering a health shock for two respondents with similar characteristics (ever smokers, persistently uninsured, similar genetic propensity to smoke, gender, etc). The only difference between the two individuals is the point in time at which they experience the shock and, hence, their exposure to the financial costs associated with it. Who responds more to the financial exposure: the high-PGS or the low-PGS type?

Methodologically, we use ordinary least squares (OLS) regression to estimate a linear probability model for smoking. Current smoking status (*Y*) is regressed on the full set of interactions between the indicators for the health shock (*shock*), being uninsured pre-65 (*uninsured*), Medicare eligibility (*post*65), and high polygenic score for smoking (*g*):

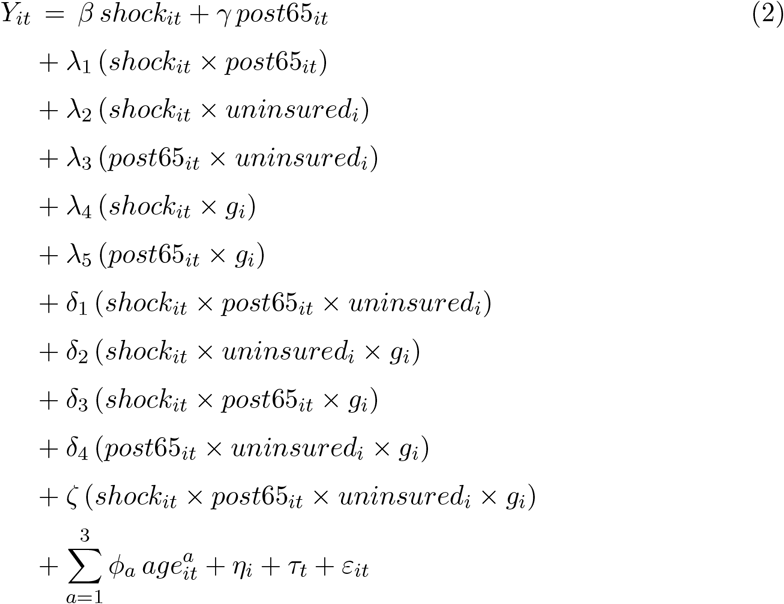

Individual fixed effects (*η_i_*) are included to control for unobserved time-invariant differences between respondents, and time fixed effects (*τ_t_*) to control for time-specific confounders. In addition, a third-degree polynomial of the respondent’s age in years at the time of the interview (*age*) is added to non-linearly control for the reduction in smoking rates shown in Figure 4. Standard errors are clustered at the individual level.

The statistical method resembles a difference-in-differences approach: focusing on the group of previously uninsured individuals, we compared the effect of a health shock for those experiencing it before or after Medicare eligibility, and who have either a low or high genetic propensity to smoke.

We are interested in 3 different types of effects for the group of previously uninsured individuals:

First, what is the marginal effect of a health shock on smoking? This effect is calculated for 4 different subgroups, given by the combinations of shock timing (pre- or post-65) and genetic risk (high or low), and shown in Figure 5. For each subgroup, the effect is comprised of the sum of all coefficient estimates from Equation (2) that applied for the given group. For example, for previously uninsured individuals who experienced the shock before the age of 65 and who had a low genetic susceptibility for smoking, the effect of the health shock is calculated by summing up the estimates for *β* and *λ*_2_. If the health shock is instead experienced after 65, the effect of the shock (for the same group of previously uninsured low-PGS individuals) is the sum of the estimates for *β*, *λ*_1_, *λ*_2_, and *δ*_1_.

**Figure 5:**
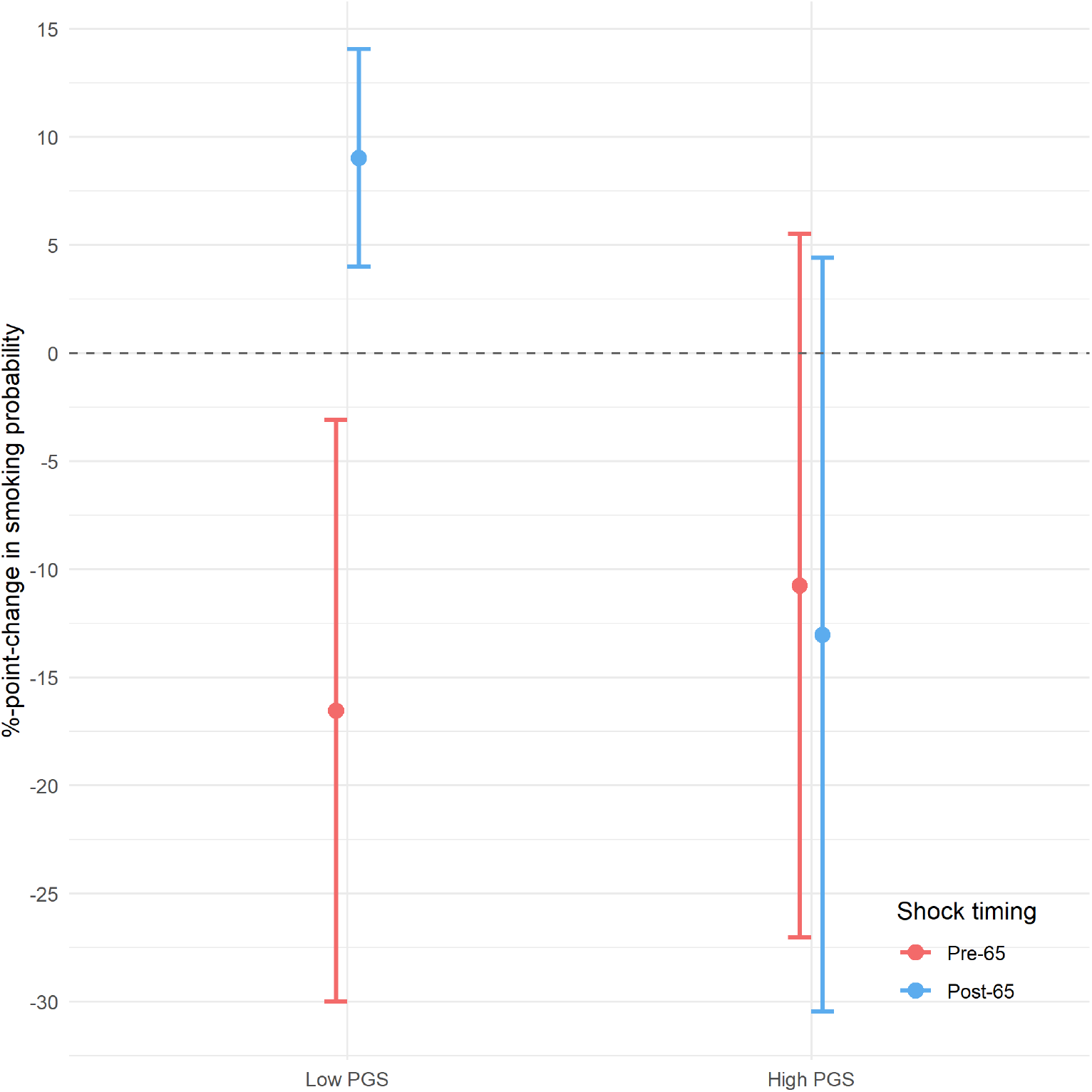
Effect of a Health Shock on the Smoking Probability in the Pre-65 Uninsured Subgroup, Stratified by Timing of the Shock and Genetic Type *Notes:* Low PGS = lowest tertile of the polygenic score distribution; high PGS = upper two tertiles of the polygenic score distribution. Pre-65: Health shock since the last survey reported at ages 60-64. Post-65: Health shock since the last survey reported at ages 67-70. Estimates and standard errors are shown in Panel A of Table 2. Effects are estimated using the coefficients in the last column of Table 3 and following the derivation described in C.5. Bars show 95% confidence intervals, standard error clustered at the individual level. *Data used*: HRS study sample, n = 5,854.

Second, what is the causal effect of Medicare eligibility (and hence a change in the financial costs associated with a health shock) on smoking? This effect is calculated using the difference between the pre-65 shock effect and the post-65 shock effect for each of the 2 genetic groups. For the low-PGS group, it is given by the sum of the estimates for *λ*_1_ and *δ*_1_. For the high-PGS group, it is given by the sum of *λ*_1_, *δ*_1_, *δ*_3_, and *ζ*.

Lastly, does the effect of health insurance vary according to individual genetic predisposition? Comparing the effect of Medicare eligibility on the post-shock smoking decision between the high- and the low-PGS groups we can assess the presence of genetic heterogeneity in moral hazard. The presence of genetic heterogeneity in moral hazard is estimated by the sum of *δ*_3_ and *ζ* in Equation (2).^8^

## 4 Results

### 4.1 Estimation Results

The effect of a health shock on the smoking probability of individuals who are uninsured before the age of 65—the subgroup of interest—for the 4 combinations of shock timing (before or after 65) and polygenic score (high or low) is shown in Figure 5 and Panel A of Table 2. Since we are controlling for individual fixed effects, these coefficients can be interpreted as the change in the probability of smoking after suffering a health shock. All of the regression coefficients are presented in the last column of Table 3.

**Table 2:** Summary of Statistical Results for the Pre-65 Uninsured Subgroup, Stratified by Timing of the Shock and Genetic Group

| **Effect of health shock on smoking probability** |  |  |
| --- | --- | --- |
|  | Low PGS | High PGS |
| Pre 65 | -0.165**<br>(0.069) | -0.108<br>(0.083) |
| Post 65 | 0.09***<br>(0.026) | -0.13<br>(0.089) |
| **Effect of health insurance on effect of health shock** |  |  |
|  | Low PGS | High PGS |
| Post 65 - Pre 65 | 0.256***<br>(0.079) | -0.023<br>(0.121) |
| **Differential effect of health insurance by genetic group** |  |  |
|  | High PGS | - low PGS |
| Post 65 - Pre 65 | -0.279*<br>(0.144) |  |

**Table 3:** Coefficients from estimating the linear probability model in equation (2) using OLS

|  | *Dependent variable:* |  |  |  |  |
| --- | --- | --- | --- | --- | --- |
|  | Smoking status |  |  |  |  |
|  | (1) | (2) | (3) | (4) | (5) |
| Health Shock | −0.027<br>(0.040) | −0.028<br>(0.040) | −0.027<br>(0.040) | −0.046*<br>(0.024) | −0.045*<br>(0.024) |
| Post-65 | −0.064***<br>(0.008) | −0.020<br>(0.012) | −0.020<br>(0.012) | −0.010<br>(0.008) | −0.009<br>(0.008) |
| Uninsured | 0.169***<br>(0.027) | 0.169***<br>(0.027) | 0.169***<br>(0.027) |  |  |
| High PGS | 0.033***<br>(0.012) | 0.033***<br>(0.012) | 0.033***<br>(0.012) |  |  |
| Shock × Post-65 | 0.008<br>(0.056) | 0.018<br>(0.056) | 0.016<br>(0.056) | 0.026<br>(0.032) | 0.025<br>(0.032) |
| Shock × Uninsured | −0.193<br>(0.184) | −0.187<br>(0.185) | −0.188<br>(0.184) | −0.118<br>(0.073) | −0.120*<br>(0.073) |
| Post-65 × Uninsured | 0.068<br>(0.048) | 0.067<br>(0.048) | 0.067<br>(0.048) | −0.052*<br>(0.031) | −0.052*<br>(0.031) |
| Shock × High PGS | 0.028<br>(0.049) | 0.029<br>(0.049) | 0.026<br>(0.049) | 0.015<br>(0.029) | 0.015<br>(0.029) |
| Post-65 × High PGS | −0.0003<br>(0.010) | −0.0005<br>(0.010) | −0.0005<br>(0.010) | −0.005<br>(0.008) | −0.005<br>(0.008) |
| Shock × Post-65 × Uninsured | 0.343<br>(0.256) | 0.336<br>(0.256) | 0.336<br>(0.255) | 0.230***<br>(0.086) | 0.230***<br>(0.085) |
| Shock × Uninsured × High PGS | 0.184<br>(0.219) | 0.182<br>(0.220) | 0.183<br>(0.219) | 0.041<br>(0.112) | 0.042<br>(0.112) |
| Shock × Post-65 × High PGS | −0.047<br>(0.068) | −0.047<br>(0.068) | −0.045<br>(0.068) | −0.078*<br>(0.042) | −0.078*<br>(0.042) |
| Post-65 × Uninsured × High PGS | −0.116*<br>(0.061) | −0.117*<br>(0.062) | −0.117*<br>(0.062) | 0.042<br>(0.036) | 0.042<br>(0.036) |
| Shock × Post-65 × Uninsured × High PGS | −0.279<br>(0.311) | −0.278<br>(0.312) | −0.278<br>(0.311) | −0.199<br>(0.151) | −0.200<br>(0.150) |
| Age |  | *Yes* | *Yes* | *Yes* | *Yes* |
| Year FE |  |  | *Yes* |  | *Yes* |
| Individual FE |  |  |  | *Yes* | *Yes* |
| Observations | 26,022 | 26,022 | 26,022 | 26,022 | 26,022 |
| R<sup>2</sup> | 0.015 | 0.017 | 0.017 | 0.823 | 0.823 |

*Notes:* Health shock = binary indicator for having suffered a cardiovascular health shock for the first time since the previous wave. Post-65 = indicator for age > 65 at the time of the interview. Uninsured = binary indicator for being persistently uninsured in every observation of the study sample before the age of 65. High PGS = indicator for being in the upper two tertiles of the polygenic score distribution. Age = controls for 3<sup>rd</sup> degree polynomial in age. FE = adding fixed effects. Robust standard errors in parentheses are clustered at the individual level. \*p<0.1; \*\*p<0.05; \*\*\*p<0.01. *Data used:* HRS study sample, n = 5,854.

Following a health shock, persistently uninsured individuals with a high polygenic score tend to reduce their smoking behavior by about 10 percentage points, irrespective of whether the health shock happens before or after the age of 65. Since the response to the health shock is the same before and after the age of Medicare eligibility, there is no evidence of moral hazard in this subgroup. With a 30.8% baseline probability of smoking, this 10 percentage point reduction is non-negligible, but it is very nosily estimated and is indistinguishable from zero.

On the contrary, the response of low polygenic score individuals differs markedly by the *timing* of the health shock: if the shock happens before the age of 65, while not covered by Medicare or another health insurance, they reduce smoking by 17 percentage points. With a baseline probability of smoking of 27%, this reduction is very sizable. On the other hand, if the health shock happens after the age of Medicare eligibility, low polygenic score individuals increase their smoking rate by 9 percentage points as compared to the rest of the population. Against the backdrop of a steady decline in smoking rates, as shown in Figure 4, this positive coefficient can be interpreted as a lower decrease in smoking for this particular sub-population.^9^

The difference in the smoking response depending on the timing of the health shock, before or after the age of 65, is summarized in Panel B of Table 2: individuals with a low polygenic score are 25.6 percentage points more likely to stop smoking if the health shock happens before the age of 65, when they are not covered by health insurance. High polygenic score individuals, on the other hand, are 2.3 percentage points more likely to do the opposite, a small and (statistically) insignificant difference.

Such marked difference in the response across the two genetic types is evidence of genetic heterogeneity in the effect of Medicare eligibility on the response to the shock. This genetic heterogeneity, a form of ‘gene-environment interaction’ (or G×E) according to behavioral geneticists,^10^ is summarized in Panel C of Table 2: low PGS individuals are 27.9 percentage points less likely than high PGS individuals to stop smoking if the health shock happens when they are uninsured.

The evidence in favor of genetic heterogeneity is remarkably stable to the inclusion of different controls and fixed effects, as shown in the different columns of Table 3. The first column does not include polynomial controls for age or any type of time or individual fixed effects; still, the sum of the estimated *δ*_3_ (*shock_it_* × *post*65_*it*_ × *g_i_*) and *ζ* (*shock_it_* × *post*65_*it*_ × *uninsured_i_* × *g_i_*) coefficients is 32.6 percentage points, close to the 27.9 estimated with age controls and time and individual fixed effects, as reported in Panel C of Table 2.

How can we interpret this evidence of genetic heterogeneity? This result is consistent with the interpretation that high polygenic score individuals are less responsive and elastic in their response to shocks and changes in the environment. Regardless of whether the health shock happens when they are covered by health insurance, and therefore whether they are financially liable for the consequences of the health shock, they reduce smoking the same amount. Individuals with a low polygenic score are instead more flexible, strategic, and reactive to the environment: they reduce their smoking only when they bear the full cost of the consequences of negative health shocks. This result is therefore consistent with genetic heterogeneity in moral hazard.

### 4.2 Other Interpretations

#### Is it really Medicare?

Other factors besides eligibility to Medicare may be at play around the age of 65, the most prominent being retirement. Such factors might affect smoking behavior after health shocks and change the interpretation of our coefficient. Our empirical result about genetic heterogeneity would still stand, but the post-65 dummy could not be considered purely as a measure of health care eligibility, and the differential reduction in smoking before 65 would not be evidence of moral hazard heterogeneity.

However, these alternative interpretations are at odds with two findings: there is no effect for those already insured before the age of 65, and there is no differential effect on income or retirement.

First, as Table 3 shows, the estimated coefficient on the shock x post-65 interaction is close to 0 and not statistically significant, suggesting no differential effect of the health shock on the probability of smoking for those individuals already covered by health insurance before the age of 65. If other factors rather than health insurance status were to cause the difference between the effects of a pre- vs. post-65 health shock in the low-PGS group, one could expect to also see an effect for those already insured before the age of 65.

Secondly, as Panels (a), (b), and (c) of Figure 6 show, there is no apparent jump in income or the probability of retirement exactly at the age of 65 in our sample. Contrasting this with the sharp change in enrollment in Medicare (Figure 3) and drop in the share of uninsured (Figure 3) suggests that the main event captured by the post-65 dummy for the sample of previously uninsured individuals is access to health coverage.

**Figure 6:**
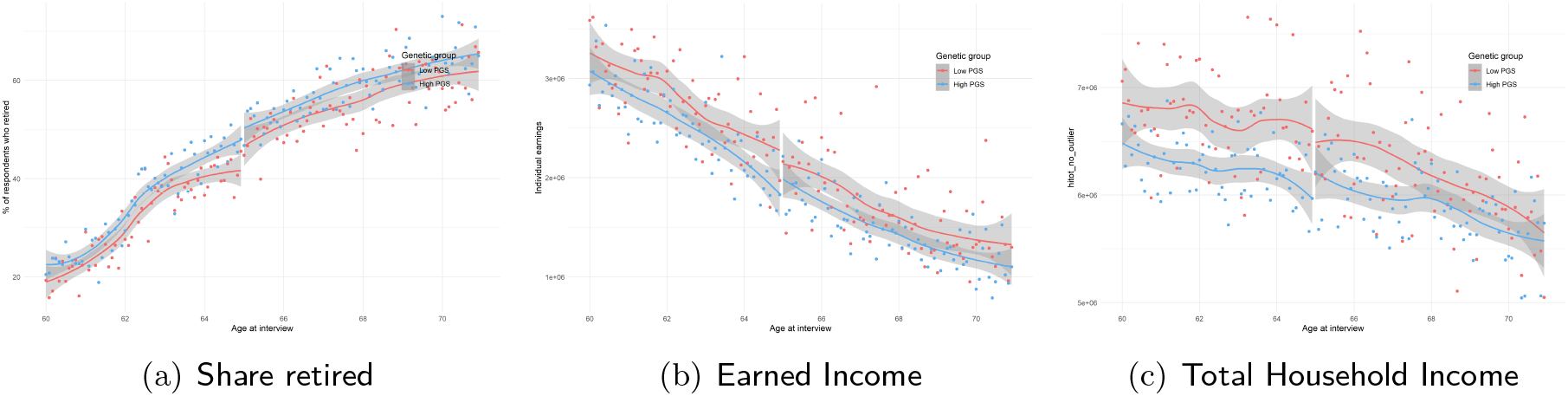
Retirement and income across the age 65 threshold by PGS. *Notes:* Self-reported retirement status, earned income, and total-household income over time. Bin-scattered plot and generalized linear smoothed correlation between age and outcome variables shown in red (low PGS = lowest tertile of polygenic score) and blue (high PGS = upper two tertiles of polygenic score). *Data used*: HRS waves 1-13, restricted to observations with age between 60 and 70 years.

Finally, as shown in Figure 8, there is no differential effect of the health shock before or after the age of 65 on the reported income or the probability of retirement of persistently uninsured individuals. The absence of any discernible effect suggests that neither income nor retirement are potential mechanisms behind the observed change in smoking responses.

#### Is it really genes?

Other individual-level characteristics, besides genetic predispositions, might be driving this heterogeneity in moral hazard. For instance, education, cognitive abilities, personality traits, or risk aversion might be a better proxy of individual level heterogeneity driving a differential response to health shocks. Given the richness of the HRS data, we can run the same analysis outlined in equation 2 but replacing *g_i_* with other individual characteristics.

None of the other individual measures that we have tried seems to be driving hetero-geneity in moral hazard, as shown in Figure 7. Specifically, we tested for years of education (highly educated people might be more knowledgeable of the insurance system), a proxy of cognitive abilities^11^ (smart people might be more strategic), risk aversion measured through hypothetical gambles on lifetime income (individual risk preferences might moderate both smoking behaviors and the response to health shocks), the Big-5 personality trait conscientiousness (conscientious people might be more likely to follow the doctor’s advice of stopping smoking after a health shock), gender (social scientist’s favorite sample split to engage in ex-post rationalizations), and individual and household income (which might be a buffer for the negative shock).^12^ For completeness, results are shown split by tertiles for continuous measures, and in two for binary measures.

**Figure 7:**
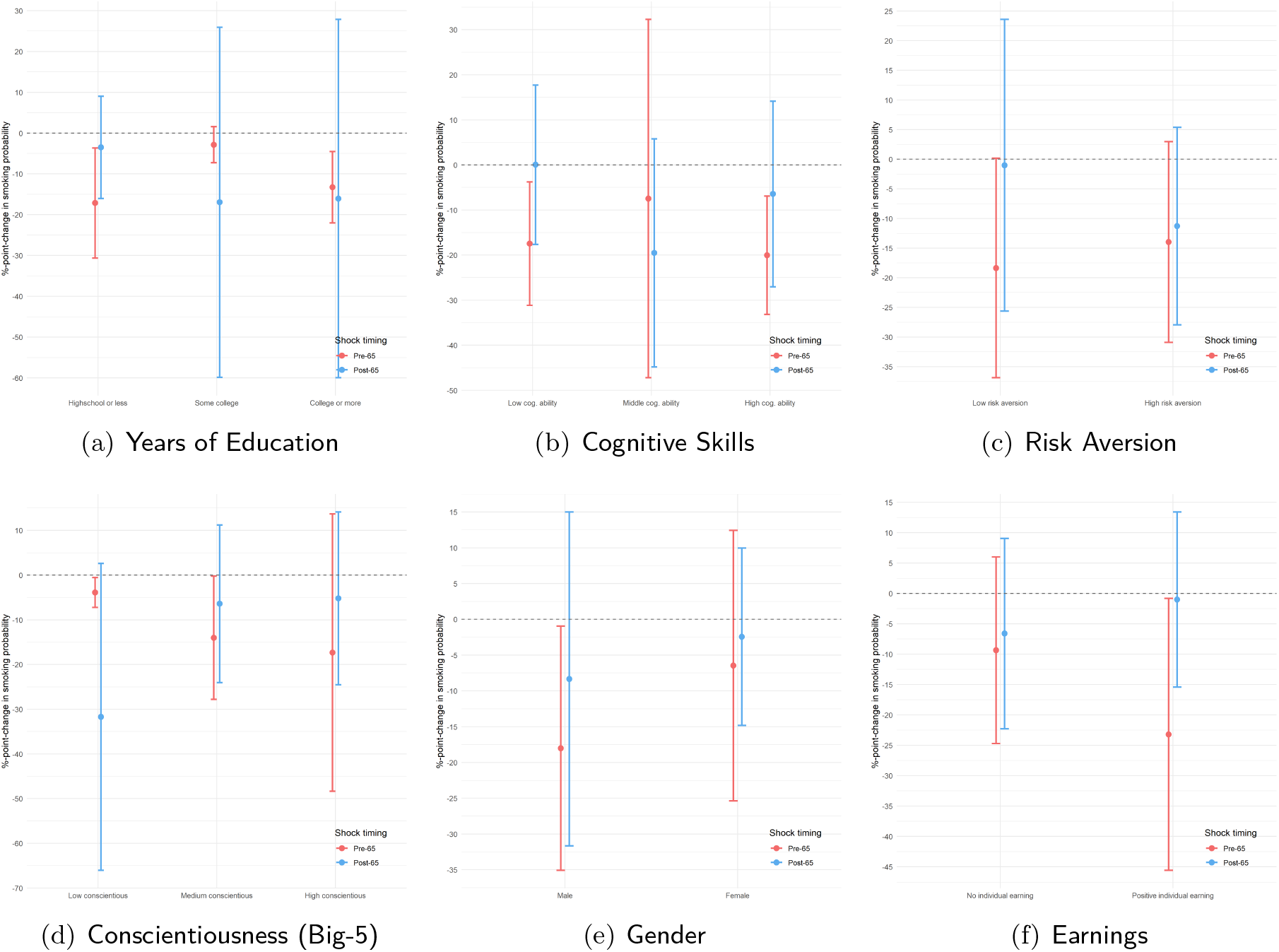
Other Individual Characteristics as Proxy for Moral Hazard Heterogeneity *Notes:* The figures report the effect of suffering a health shock on the probability of smoking for the pre-65 uninsured subgroup, stratified by timing of the shock (before and after the age of 65) and different measures of individual characteristics. Effects are estimated using a combination of the coefficients from equation 2 where *g_i_* is replaced by the different individual characteristics reported in the sub-figure captions, following the derivation described in C.5. Pre-65: Health shock since the last survey reported at ages 60-64. Post-65: Health shock since the last survey reported at ages 67-70. Bars show 95% confidence intervals, standard error clustered at the individual level. *Data used*: HRS study sample, n = 5,854.

#### Is this driven by confounders?

One might worry that the differential change in smoking behavior might be driven by other confounding factors that coincidentally happen around the age of 65, or are triggered by the health shock. For example, suffering a heart attack might reduce people’s income, induce them to retire, change their marital status, increase their out-of-pocket medical spending, or shorten their life expectancy. Any of these changes happening differentially for people with high or a low polygenic score, and before or after the age of 65, could invalidate our results or at least our interpretation of the effect as evidence of moral hazard. To test the plausibility of these concerns, we estimate equation 2 again including each of these potential confounders as an outcome variable *Y_it_*, as suggested by Pei, Pischke and Schwandt (2018).

None of these potential confounders seems to be driving our results, as shown in Figure 8. If anything, the symmetric results for out-of-pocket medical expenditure shown in panel (e) suggest that the size of the health shock is comparable for both high and low PGS individuals: both report an increase of about 2% in out-of-pocket medical expenditure if the shock happens before the age of 65 (when uninsured) and a change smaller than 0.3% if the shock happened after for low PGS, and a reduction of about 1% for those with high PGS (and not statistically significant for neither of these last two estimators).

**Figure 8:**
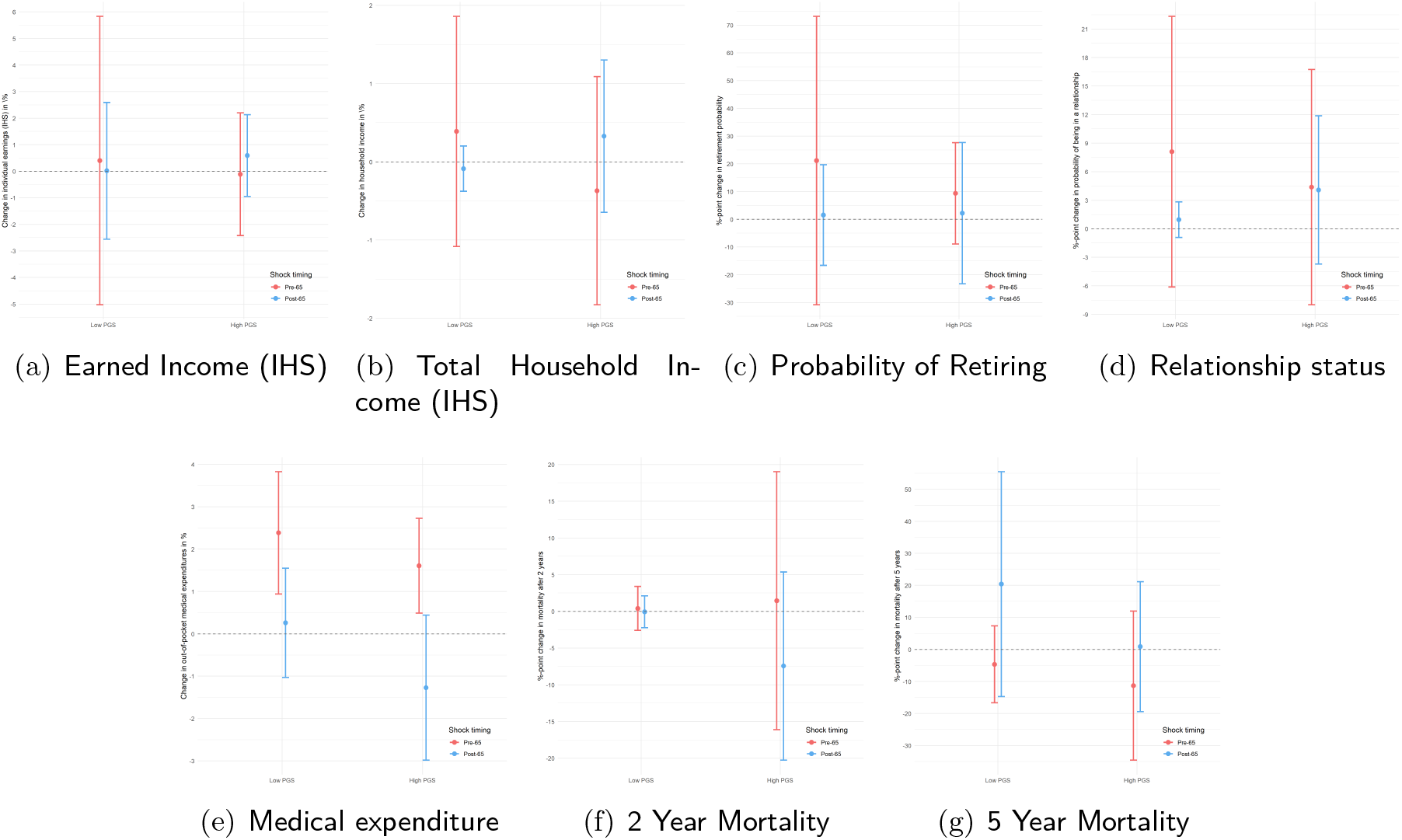
Testing for Potential Confounders *Notes:* The figures report the effect of suffering a health shock for the pre-65 uninsured subgroup, stratified by timing of the shock (before and after the age of 65) and having a high or low polygenic score, on various possible confounders. Effects are estimated using a combination of the coefficients from equation 2 where the outcome *Y_it_* is replaced by the different confounders reported in the sub-figure captions, following the derivation described in C.5. Low PGS = lowest tertile of the polygenic score distribution; high PGS = upper two tertiles of the polygenic score distribution. IHS: inverse hyperbolic sine (similar to log). Pre-65: Health shock since the last survey reported at ages 60-64. Post-65: Health shock since the last survey reported at ages 67-70. Estimates and standard errors are shown in Panel A of Table 2. Effects are estimated using the coefficients in the last column of Table 3 and following the derivation described in C.5. Bars show 95% confidence intervals, standard error clustered at the individual level. *Data used*: HRS study sample, n = 5,854.

### 4.3 Robustness checks

These results are robust to changes in the definition of the high-PGS indicator (having a PGS above the median, using a linear PGS, using an older GWAS for the weights) and the definition of the pre-65 uninsured status indicator (uninsured in 33% or 66% of all pre-65 observations instead of 100%). Relaxing the definition of the uninsured indicator to include respondents uninsured in a minimum of 33% of pre-65 observations leaves the directions of the effects unchanged, but the magnitudes are smaller and statistical significance is lost. Furthermore, the findings of this study do not depend on the exclusion of HRS respondents for whom Medicare eligibility status at the time of the shock is unknown (when health shocks are reported at ages 65 or 66). Estimation results for all robustness checks are shown in Section D.3 in the Appendix.

## 5 Discussion

In this study, experiencing a cardiovascular health shock is associated with a significant reduction in the smoking probability of uninsured 60-to 64-year-old individuals with a low index of genetic propensity to smoke. Medicare eligibility after age 65 (and hence a lower exposure to the financial costs of illness) fully neutralized this cessation effect, indicating the presence of moral hazard caused by insurance coverage. For individuals with a high genetic propensity to smoke, experiencing a health shock does not significantly affect the smoking probability, irrespective of whether the shock is at a time of high or low exposure to the financial costs of illness.

The effect of health shocks on changes in smoking behavior, as well as the underlying mechanisms, have been addressed before (Clark and Etilé, 2002; Falba, 2005; Keenan, 2009; Khwaja, Sloan and Chung, 2006*a*,*b*; Marti and Richards, 2017; Richards and Marti, 2014; Smith et al., 2001; Sundmacher, 2012; Wray et al., 1998). Previous empirical work has found strong evidence of an increase in smoking cessation after a health shock (Clark and Etilé, 2002; Falba, 2005; Keenan, 2009; Marti and Richards, 2017; Richards and Marti, 2014; Sundmacher, 2012; Wray et al., 1998). The mechanism that has received a lot of attention in earlier research is a changed perception of personal health risks and survival probability, motivating the individual to reduce tobacco consumption to improve future health (Clark and Etilé, 2002; Khwaja, Sloan and Chung, 2006*a*,*b*; Smith et al., 2001).

Marti and Richards (2017); Richards and Marti (2014) have highlighted the role that financial costs associated with health shocks, as opposed to only the health considerations, can play in determining the post-shock smoking decision. In individuals with high financial risk exposure, health shocks may bring about significant out-of-pocket medical costs. At the same time, the financial consequences of smoking-related illness are likely more complex and opaque to the individual than the health-related consequences of smoking.

The mechanism presented in these studies suggests that through improving their grasp on the financial cost of smoking, the increase in out-of-pocket health care costs following the health shock can serve as an impetus for smoking cessation. Using the same design that we follow in this analysis (the age-based eligibility threshold for the Medicare program), a recent study has provided robust evidence for this mechanism (Marti and Richards, 2017). Without investigating potential heterogeneity between genetic groups, they found that, on average, moral hazard is present, and Medicare eligibility reduced the cessation effect that a cardiovascular health shock had for uninsured individuals.

Our analysis shows that this average effect is likely driven by individuals with a low genetic predisposition for smoking only. The effect cannot be observed for the subgroup of individuals with a high genetic predisposition for smoking, suggesting that genetic makeup can act as a constraint and limit the extent to which incentives for behavior change are translated into actual behavior change.

The question of how insurance policies can interact with genetic predispositions for risky health behaviors is interesting from a policy perspective for several reasons. The finding that insurance affected the post-shock smoking decision only in part of the sample suggests that the considered type of moral hazard in health insurance, which causes excess smoking among an already sick population, is less prevalent than an initial inspection may suggest. This insight may alleviate one possible concern against universal public health care coverage (Einav and Finkelstein, 2018; Mendoza, 2016).

On a broader level, awareness of the possibility of interactions between genetics and health insurance is important for understanding how the health insurance system can increase or reduce genetically induced health inequalities in a society. By highlighting the role of genetic influences on the propensity for and ability to quit unhealthy behaviors, this study also points to possible limitations on the extent to which insurance can cause health behavior changes through financial incentives.

Genetically determined limitations on the effectiveness of financial incentives are important to consider not just when evaluating the effectiveness of health insurance policies, but also the fairness therein. With recent technological advances, for example in the field of wearable tech, health insurers are discovering more and more possibilities for monitoring health behaviors. This information is increasingly used for pricing, with the explicit goal of motivating behavior change through financial incentives in the form of lower insurance premiums or deductibles (Olson, 2014; Young, 2017).

In light of this development, the possibility that genetic predisposition prevents individuals from changing health behaviors, despite strong incentives for doing so, is becoming increasingly relevant. By emphasizing the correlation between genetic risk and an inability to quit unhealthy behaviors, the findings in this study may raise the following question: To what extent do insurance policies that price differentiate based on health behaviors ultimately also discriminate based on genetics? Under current US legislation, the 2008 Genetic Information Nondiscrimination Act prohibits health insurers from discriminating based on explicit genetic information. Not wanting to punish those who are disadvantaged in terms of their genetic makeup is one of the motivations behind this legislation. To create a health insurance system that can incentivize healthy behaviors and simultaneously reflect society’s perception of fairness, it will be important for future research to further enhance our understanding of how genetics and health insurance interact and jointly affect health behaviors.

### 5.1 Limitations

This study has a few limitations. First, the results hinge on a small number of respondents who suffer a health shock around the age of 65. Although the total sample size used in the regression is greater than 5 thousand individuals, the number of people in each cell (i.e. those who suffer a health shock, before or after the age of 65, and have a certain polygenic score) is between 110 and 274.^13^ We encourage and welcome a replication of our results, preferably in a within-family design which can better account for the thorny issue of ancestry and population structure, but we are unaware of an existing dataset that contains all of the necessary information about genetic predispositions, health insurance coverage, and smoking behaviors.

Second, the analysis focused on the short-run smoking response to a health shock, and only considered changes in the extensive margin—i.e., changes between smoking and not smoking. From a public health perspective, interactions of genetics with both the long-term persistence of the behavior change as well as behavior changes along the intensive margin of smoking—i.e., changes in the number of cigarettes smoked or the intention to quit smoking—may be of interest.

Third, a few caveats regarding the internal validity of the study. This study relied on self-reported information regarding smoking behavior, health diagnoses, and insurance status. If participants of different ages and different genotypes differentially misreported their smoking status, for example, the estimated effect of Medicare eligibility on the smoking response to a health shock could be biased. We have found no published evidence that misreporting might be associated with individual genotypes. It is also possible that those identified as continuously uninsured before the age of 65 had insured spells in between the biennial HRS interviews. These unmeasured episodes with coverage may have biased the results towards not finding a significant effect of Medicare eligibility. Finally, this study may also suffer from survivorship bias and attrition. Because DNA collection and genotyping took place relatively late during the HRS (starting in 2006), it is possible that study participants with very high genetic susceptibility for smoking, and hence particularly unhealthy smoking habits, passed away before the DNA collection and are systematically excluded from the study population. Within the study population, participants with the highest genetic propensity to smoke may have been less likely to reach the age of Medicare eligibility, hence leading to relatively lower genetic risk in the group that experienced health shocks post-65. Both problems would likely have lead to an underestimation of the difference between lowand high-PGS individuals.

Lastly, a few words of caution on the external validity of the study. The transition from being uninsured to being eligible for Medicare at age 65 is a setting that is very specific to the health care system in the US. While Medicare can generally be seen as just one example of universal health care coverage, it only applies to a specific age group, and it is not clear whether smoking behaviors in response to a health shock in this age group are representative for all ages in the population. Another possible concern may be that Medicare eligibility does not necessarily translate into Medicare coverage, as take-up rates are not at 100%. As Figure 3 shows, however, the fraction of eligible HRS respondents in the study sample who are actually enrolled in Medicare is high—approximately 90% at age 65, increasing to 98% at age 70. Additionally, take-up patterns do not differ much between the genetic groups. Table 14 in the Appendix shows that using actual Medicare enrollment status rather than Medicare eligibility status in the empirical analysis does not change the results.

## 6 Conclusions

Medicare eligibility significantly lowered the probability of smoking cessation after a health shock in individuals aged between 60 and 70 years who are uninsured before the age of 65 and have a low genetic predisposition for smoking. Health insurance can plausibly affect the smoking response to a health shock by lowering the financial risk associated with the shock, and thereby eroding additional incentives for behavior change. This change in behavior following Medicare eligibility is indicative of moral hazard and is not observable among individuals with a high genetic predisposition for smoking. The differential effect of Medicare eligibility for the two genetic groups suggests that biological constraints can overpower both health-related and financial incentives for smoking cessation, and provides a readily available measure of heterogeneity in moral hazard. Heterogeneity in moral hazard can be used to enrich economic models of health behavior^14^ and our understanding of how individual biological characteristics can influence decision-making.

Building on previous work analyzing the interplay between genes and exogenous environmental changes (Barcellos, Carvalho and Turley, 2018; Fletcher, 2012; Schmitz and Conley, 2017*a*), this study provides a contribution to the centennial debate about nature *and* nurture (Galton, 1874; Haldane, 1946; Kong et al., 2018; Lundborg and Stenberg, 2010; Mulcaster, 1582), casting further doubts about genetic determinism. The influence of genetic variants on our choices and outcomes is modulated by the environment around us, just as the response to environmental shocks is filtered through the prism of our genetic predispositions.

Our results show that genetic factors can influence health decisions and strategic behaviors, and therefore should be taken into consideration when evaluating the effectiveness and fairness of different policies, such as health insurance. Fairness considerations should take into account the fact that genetic endowments are passed down from one generation to the next, are fixed at conception, and cannot be changed by an individual’s choices or effort, raising questions of deservedness, merit, and luck (Harden, In Press; Kweon et al., 2020; Pereira, 2021). Efficiency considerations should consider that genetic endowments are usually unobserved to the individual, the insurance companies, and the government. This unobservability raises important regulatory questions at the intersection between health and information economics: who, if anyone, should have this information? How should the information be provided? How will this information affect demand and supply? Should private contracts or public policies take this information into account? The recent rise of direct-to-consumer genetic testing services might force this discussion into the public debate sooner than expected.

More generally, future studies should build off of the idea of leveraging existing indices of genetic predispositions to provide biological measures of heterogeneity in human behaviors. This can enrich economic models and empirical studies, shedding new light on fundamental economic parameters.

## A Appendix

## B Genetics for Economists

The human genome consists of over 3 billion base pairs (6 billion bases) in each cell nucleus, with four possible bases: adenine (A), thymine (T), guanine (G), and cytosine (C).^15^ Comparing any two unrelated human beings, over 99% of their genome will be identical. The remaining < 1% differs between individuals, with a Single Nucleotide Polymorphism (or SNP, pronounced *snip*) being the most common form of genetic variation. A SNP is a single base-pair substitution at a particular location (locus) on the human genome.

To identify genetic variants that are associated with a particular trait of interest, such as coronary heart disease, so-called Genome-Wide Association Studies (GWAS) relate each SNP to the trait in a hypothesis free-approach. Stringent p-values are then used to identify SNPs that are robustly associated with the trait of interest, and replication is performed in other, independent samples. Only SNPs that have consistent associations across the different samples are interpreted as robust. However, this does not guarantee that individual SNPs have large effect sizes. Most human complex traits are polygenic, meaning that they are affected by many SNPs, each with a very small effect size. To increase the predictive power of SNPs, they can then be aggregated into so-called polygenic scores (PGS), defined as:

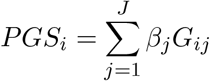

where *G_ij_* is SNP *j* of individual *i*, and *β_j_* is the effect size of that SNP, obtained from an independent Genome-Wide Association Study. GWAS sample sizes have grown substantially in recent years, meaning that (a) SNPs with very small effects are more likely to be identified, and (b) that the effect sizes are estimated with increased precision. Indeed, we have seen large improvements in genetic prediction, with initial PGS being able to explain less than 1% of the variation in the trait of interest, to more recent ones explaining up to 11-13% of the variation in educational attainment just by increasing the sample size of the discovery sample (see e.g., Lee et al., 2018). PGS have been shown to be powerful tools to identify patients with increased risk for coronary artery disease, atrial fibrillation, type 2 diabetes, inflammatory bowel disease, and breast cancer. Indeed, Khera et al. (2018) propose the use of polygenic prediction in clinical care.

## C Methods

## C.1 DNA Extraction and Genotyping

In 2006, the Health and Retirement Study (HRS) introduced enhanced face-to-face interviews (EFTFs), which expanded the core interview with measures of physical function, blood-based biomarkers, and DNA samples. Sample selection for the EFTFs was conducted as follows: A random 50% of the 2006 sample was preselected for an EFTF, and the other half was selected in 2008. A new cohort of households was added to the HRS in 2010. Of these new households, a random 50% was selected for EFTF data collection in 2010, while the other half was selected in 2012. The households selected for EFTFs in 2012 are not yet included in the polygenic scores data used in this study. In 2006, saliva collection was conducted using a mouthwash collection method. From 2008 onwards, the Oragene DNA Collection Kit (OG-250) was used.

Genotype data was obtained for over 15,000 HRS participants. Genotyping was conducted by the Center for Inherited Disease Research (CIDR) in 2011, 2012, and 2015, using the Illumina HumanOmni 2.5 BeadChips (HumanOmni2.4-4v1, HumanOmni2.5-8v1). Approximately 2.4 million single nucleotide polymorphisms (SNPs) were measured. Of the roughly 1.9 million genotyped SNPs that passed quality control, 21 million SNPs were imputed using the 1000 Genomes Reference Panels (phase 3, version 5). More details on genotyping and imputation can be found in the official HRS Documentation Report.

## C.2 Study Sample

To improve replicability of the results, we mostly use data from the publicly available RAND HRS file (version P)(, n.d.*a*)—an easy-to-use longitudinal data set based on the HRS data—as well as the publicly available initial release of the HRS polygenic scores data (, n.d.*b*).

## C.2.1 Reshaping, Merging, and Sample Restrictions

This section describes how the study sample was constructed from the RAND HRS version P data file. First, the data file was reshaped from wide to long format, with each observation corresponding to a respondent-wave entry. Second, polygenic risk scores (PGSs) for the HRS phenotype “smoking initiation” (referred to as “regular smoking” in this study, for clarity purposes) from the initial HRS PGS data release, using genetic data from 2006 to 2010, were merged for the 9,991 genotyped individuals of European ancestry. The following shows the list of restrictions that were then imposed to arrive at the study sample used in the main analysis. Regarding notation, note that VARIABLE refers to the long-format version of the variables that were called R1VARIABLE to R12VARIABLE in the RAND HRS data file. From the reshaped data file, the study sample was reached by carrying out the following steps (in this exact order):

1. Drop observations with an age (AGEY E; see Section C.3.3) below 60 or above 70 years
2. Drop individuals with only 1 observation
3. Drop observations with missing values for the PGS for smoking (PGS EvrSmk TAG10; see Section C.3.5), the self-constructed health shock indicator (Section C.3.2), or the current smoking status (SMOKEN; see Section C.3.1)
4. Drop individuals who in their first observation (the baseline) reported never having smoked (SMOKEV equal to 0; see Section C.2.2)
5. Drop individuals with missing values for the self-constructed pre-65 uninsured status indicator (Section C.3.4)
6. Drop individuals who reported a health shock (self-constructed health shock indicator equal to 1; see Section C.3.2) when interviewed at ages 65 or 66

## C.2.2 Ever-Smoker Status

“Ever-smoker” refers to the RAND variable SMOKEV, which indicates whether the respondent has ever smoked cigarettes. Ever smoking means having smoked more than 100 cigarettes throughout one’s life, not including pipes or cigars. This is consistent with the Centers for Disease Control classification of the term “ever-smoker.”Jamal et al. (2016) The ever-smoked question was usually only asked at the respondent’s first interview and then carried forward for subsequent waves. For details on the survey questions and recodings for missings into yes/no answers, see the publicly available official RAND HRS documentation.

## C.3 Outcome and Exposure Variables

## C.3.1 Smoking Status

Current smoking status refers to the RAND variable SMOKEN, which indicates whether the respondent smokes at the time of the interview. The survey question about current smoking status was only asked for respondents who answered yes to being ever-smokers (having smoked more than 100 cigarettes in their lifetime). For details on the survey questions and recodings for missings into yes/no answers, see the RAND HRS documentation.

## C.3.2 Health Shocks

The health shock indicator was defined using the RAND variables HEARTE and STROKE. The variable HEARTE indicates whether or not a doctor has ever told the respondent that he/she had 1 of the following conditions:

1. Heart attack
2. Coronary heart disease
3. Angina
4. Congestive heart failure
5. Other heart problems

The variable STROKE indicates whether or not a doctor has ever told the respondent that he/she had 1 of the following conditions:

1. Stroke
2. Transient ischemic attack

For details on the survey questions and the construction of these variables, see the RAND HRS documentation.

The health shock indicator used in this analysis was then defined as follows: It was set to 1 in a given wave if the lagged values of both HEARTE and STROKE were equal to 0, and if either one of the current values (or both) was equal to 1.

## C.3.3 Medicare Eligibility Status

The Medicare eligibility status indicator was defined using the RAND variable AGEY E. This variable indicates the respondent’s age in years at the end of the HRS interview in a given wave. For details on the construction of this variable, see the RAND HRS documentation.

## C.3.4 Pre-65 Uninsured Status

The pre-65 uninsured status indicator was defined using the RAND variables HIGOV, COVR, COVS, and HIOTHP. The variable HIGOV indicates whether the respondent was covered by any government health insurance program. COVR indicates whether the respondent was covered by health insurance from his/her current or previous employer. COVS indicates whether the respondent was covered by his/her spouse’s employer. HIOTHP indicates whether the respondent was covered by any health insurance other than the government, employer-provided, or long-term care insurance. For details on the survey questions and the construction of these variables, see the RAND HRS documentation.

The pre-65 uninsured status indicator used in this analysis was then defined as follows: First, we defined a wave-specific uninsured indicator, which was set to 1 in a given wave if the values of HIGOV, COVR, COVS, and HIOTHP were all equal to 0. Then, the pre-65 uninsured status indicator was set to 1 for respondents whose wave-specific uninsured indicator was 1 in 100% of all pre-65 observations (i.e., where AGEY E < 65).

## C.3.5 Polygenic Risk for Regular Smoking

The high-PGS indicator was defined using the PGS for the phenotype “smoking initiation” (referred to as “regular smoking” in this study, for clarity purposes) using **all of the genetic variants**(SNPs).

The PGS was calculated using the effect sizes estimated in a genome-wide association study (GWAS) meta-analysis Liu et al. (2019) conducted by the GWAS and Sequencing Consortium of Alcohol and Nicotine use (GSCAN). The phenotype “smoking initiation” studied in this GWAS was defined as ever versus never having been a regular smoker, where regular smokers were individuals who reported having smoked ≥ 100 cigarettes throughout their life.

The PGSs used these GWAS-estimated effect sizes for all SNPs that overlapped between the HRS genetic database and the GWAS meta-analysis, without accounting for linkage disequilibrium between SNPs or considering P-value thresholds. Scores were calculated according to Equation (1) in the main text using the software packages PRSice and PLINK.

The high-PGS indicator used in this analysis was then defined as follows: It was set to 1 for individuals with a PGS above the lowest tertile, and to 0 for individuals with a PGS below or equal to the lowest tertile. The two upper-tertiles of the PGS distribution were combined to improve statistical power and simplify the exposition. Initial results using an indicator for each tertile of the distribution, displayed in Appendix Section C.5.2, show that the results for the two upper-tertiles of the PGS distribution are very similar to each other.

The polygenic score is predictive of smoking behavior, as expected and displayed in Figure 1, but not only. As shown in Appendix Figure 9—which displays the coefficients of simple OLS regressions of several outcomes on the linear PGS controlling for age, gender, and the 10 principal components of the genomewide matrix—the PGS is also predictive of other unfavorable outcomes: younger age at first birth as well as age started smoking, lower cognitive skills, perseverance, years of education, wealth, income, health rating, higher depressive symptoms, anxiety, non-cancer illnesses, drinking behavior. Reassuringly, the correlations with retirement, medications taken, and mortality are positive but extremely small and not distinguishable from zero.

**Figure 9:**
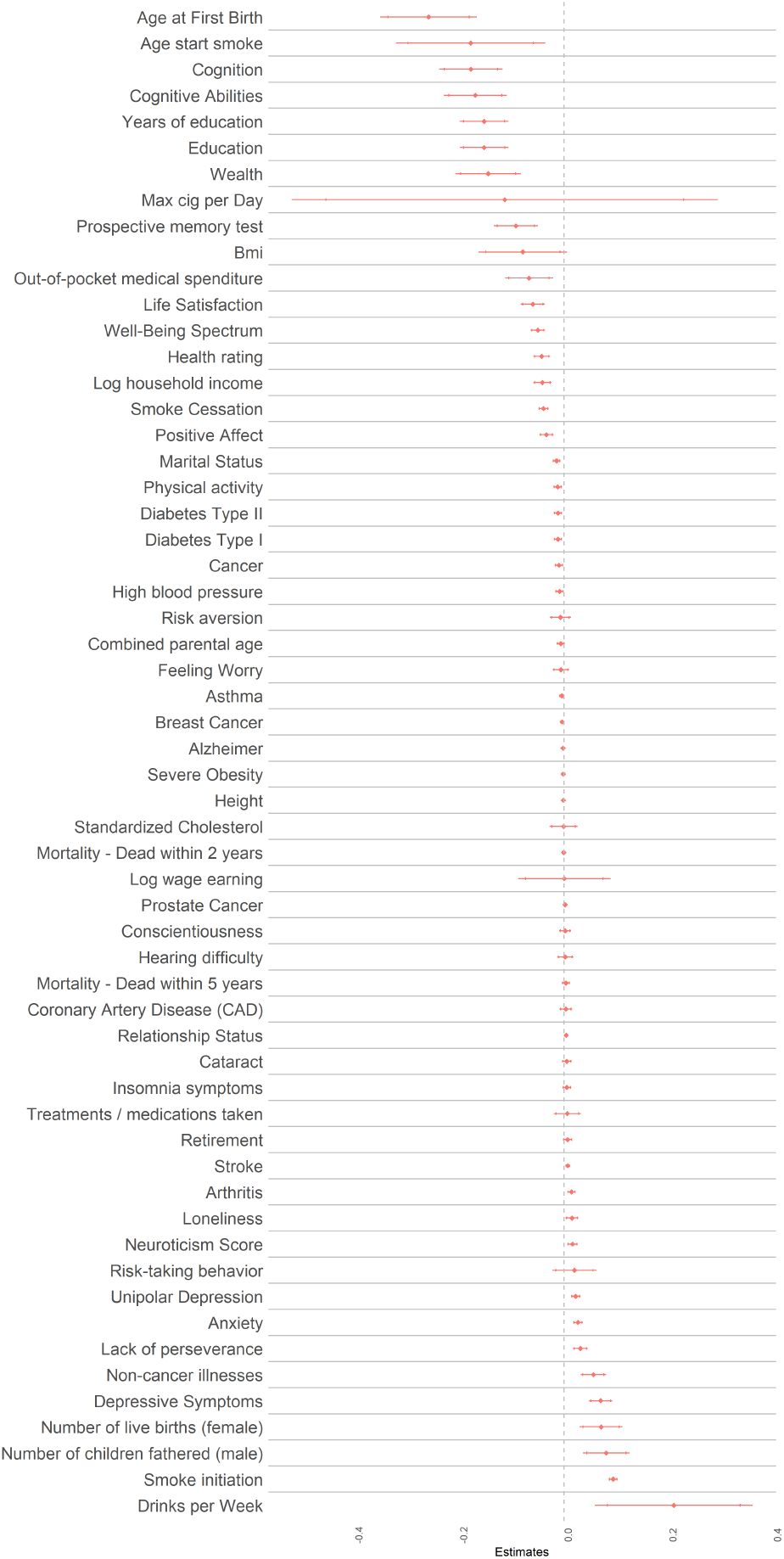
PGS distribution and correlation with smoking behavior. *Notes:* Plot of estimated coefficients associated with the PGS (entered linearly) from several OLS regressions of the different outcomes displayed on the y-axis on the PGS, age, age squared, age cubed, sex, and the first 10 principal components of the genomewide matrix. *Data used*: HRS waves 1-13, restricted to observations with age between 60 and 70 years.

As shown in Figure 10, the PGS is mildly correlated with gender and almost uncorrelated with the probability of suffering from a health shock.

**Figure 10:**
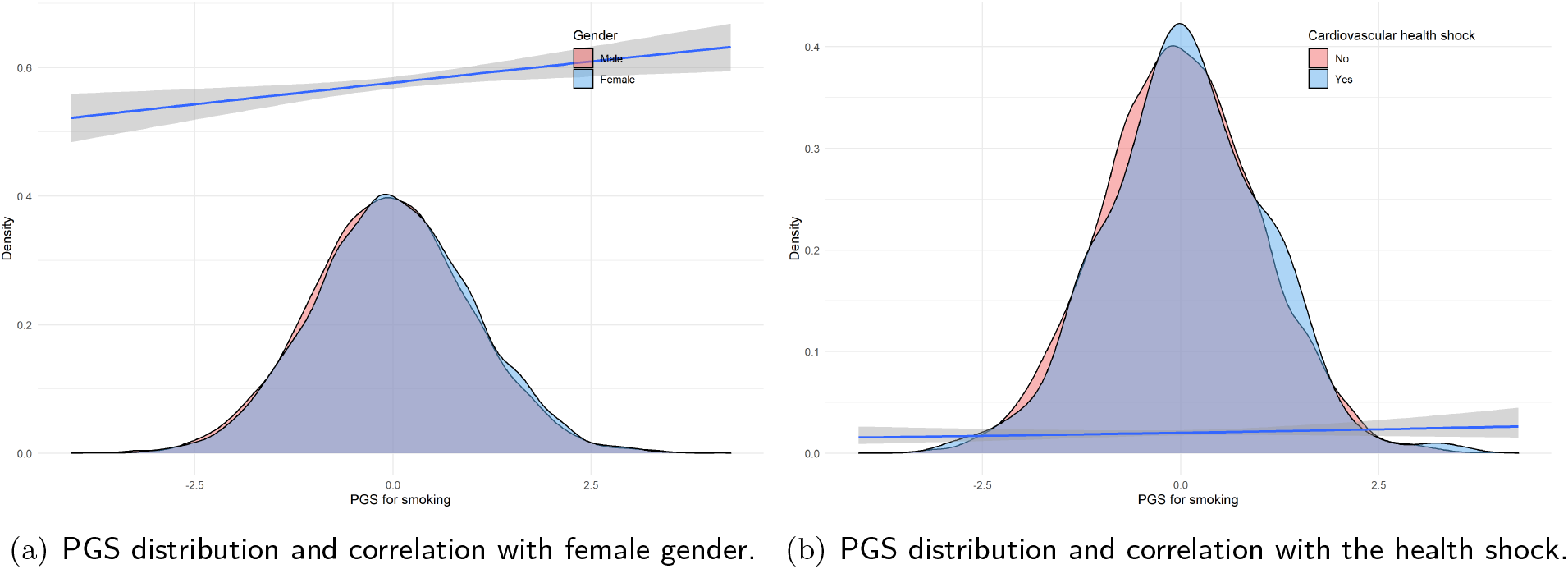
PGS for Regular Smoking and Smoking Behavior in the HRS Data *Notes:* Distribution of Polygenic Score (PGS) for baseline smokers (blue) and non-smokers (red). Generalized linear smoothed correlation between current smoking and PGS shown in the blue line (with 95% confidence intervals in shaded grey area). *Data used*: HRS waves 1-13, restricted to observations with age between 60 and 70 years.

## C.4 Statistical Analysis

## C.4.1 Age Pattern of Health Shock Incidence

With the data used in this study, it was not possible to narrow down the exact timing of a health shock to more than the between-survey 2-year window. Therefore, the probability of having a health shock at a specific age could not be determined. What could be said from this data about the age at the health shock is that for all shocks reported at ages 64 or below, the shocks must have occurred before the age of 65. Similarly, for all health shocks reported at ages 67 or above, the shocks must have occurred after the age of 65. For shocks reported at interview ages 65 or 66, it could not be determined whether the shock occurred before or after age 65 (as interviews were conducted biennially).

Figure 11 visualizes the fraction of HRS respondents who reported experiencing a health shock since the last survey wave at a given interview age, stratified both by genetic group and by gender. By visual inspection, there seems to be a positive trend in the fraction of respondents reporting health events with age, with frequent deviations but no obvious jump between 64 and 67.

To formally test for a jump or change in trend in the incidence of health shocks around the age of 65, a segmented regression approach was used. Specifically, we tested if the change between the percentage of respondents who reported a health shock at age 64 and the percentage of respondents who reported a health shock at age 67 was larger than what could be explained by a linear age trend. For this test, all observations where respondents were aged 65 or 66 were excluded. For all remaining observations, the binary health shock indicator (*shock*) was regressed on the age variable (*age*), a post-age-67 indicator variable (*post*67), and a post-age-67 trend (*post*67*slope*):

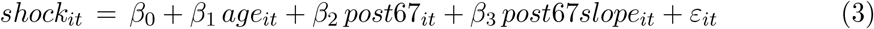

**Figure 11:**
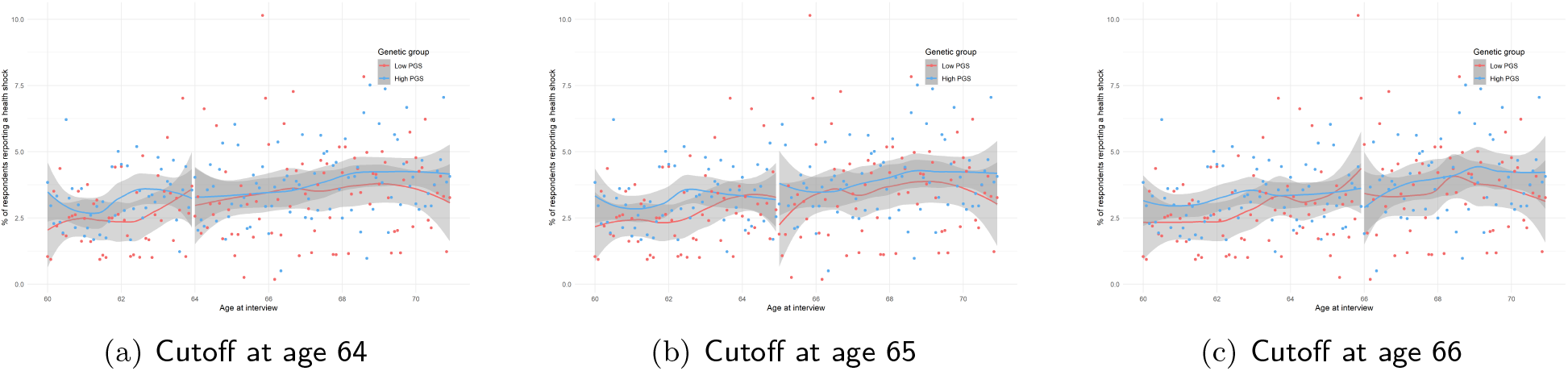
Percentage of Reported Health Shocks by Age Self-reported indicator of having been diagnosed for the first time with a cardiovascular condition since the last HRS survey. Age refers to the time of the survey, not the time of the health shock, which is unknown up to a 2-year windows, since HRS surveys are bi annual. Bin-scattered plot and generalized linear smoothed correlation between age at interview and cardiovascular health shock shown in red (low PGS) and blue (high PGS). Linear smoothed correlation estimated separately to the left and to the right of age cutoff: (a) age 64; (b) age 65; (c) age 66. *Data used*: HRS waves 1-13, restricted to observations with age between 60 and 70 years.

The post-age-67 indicator variable was defined to take the value 1 if a respondent was aged 67 or older at the time of the HRS interview. Therefore, it guaranteed that any potential health shocks were experienced after the age of 65. The post-age-67 slope variable was a continuous variable coded 0 up to and including age 67, and increased sequentially from 1 thereafter. *β*_1_ captured the general age trend in the probability of reporting a health shock; *β*_2_ estimated the jump in the report of health shocks at age 67; *β*_3_ reflected changes in the age trend of reported health shocks after age 67.

The linear probability model in Equation (3) was estimated using ordinary least squares (OLS) regression for (i) the study sample (HRS waves 1-12, restricted to observations with age between 60 and 70 years, and additionally excluding all observations with ages 65 or 66), (ii) both genetic groups separately, and (iii) both men and women separately. Estimation results are shown in Table 4. Across all groups, there were no statistically significant jumps for health shocks reported at age 67 compared to age 64 (accounting for a linear age trend). Similarly, the age trend was not significantly different after age 67 than before. In the study sample and in the low-PGS group, the increase with age in the probability of reporting a health shock was statistically significant.

**Table 4:** Coefficients from Estimating the Linear Probability Model in Equation (3) Using OLS

|  | *Dependent variable:* |  |  |  |  |
| --- | --- | --- | --- | --- | --- |
|  | Probability of shock |  |  |  |  |
|  | All | Low PGS | High PGS | Male | Female |
|  | (1) | (2) | (3) | (4) | (5) |
| Age | 0.002*<br>(0.001) | 0.003*<br>(0.001) | 0.001<br>(0.001) | 0.001<br>(0.001) | 0.002*<br>(0.001) |
| Post-67 dummy | −0.001<br>(0.005) | −0.007<br>(0.008) | 0.002<br>(0.006) | −0.001<br>(0.008) | −0.001<br>(0.006) |
| Post-67 slope | −0.0005<br>(0.001) | −0.002<br>(0.002) | 0.0003<br>(0.002) | 0.001<br>(0.002) | −0.002<br>(0.002) |
| Constant | −0.068<br>(0.049) | −0.134<br>(0.082) | −0.036<br>(0.060) | −0.046<br>(0.079) | −0.082<br>(0.061) |
| Observations | 39,185 | 13,076 | 26,109 | 16,648 | 22,537 |

*Note:*
\*p<0.1; \*\*p<0.05; \*\*\*p<0.01
Low PGS = lowest tertile of the polygenic score distribution; high PGS = upper two tertiles of the polygenic score distribution.
*Data used:* HRS waves 1-13, restricted to observations with age between 60 and 70 years and non-missing smoking status, and additionally excluding all observations with ages 65 or 66.

## C.5 Derivation of the effects from the OLS coefficients

## C.5.1 OLS estimation: high and low PGS

Current smoking status (*Y*) is regressed on the full set of interactions between the indicators for the health shock (*shock*), being uninsured pre-65 (*uninsured*), Medicare eligibility (*post*65), and high polygenic risk for smoking (*g*):

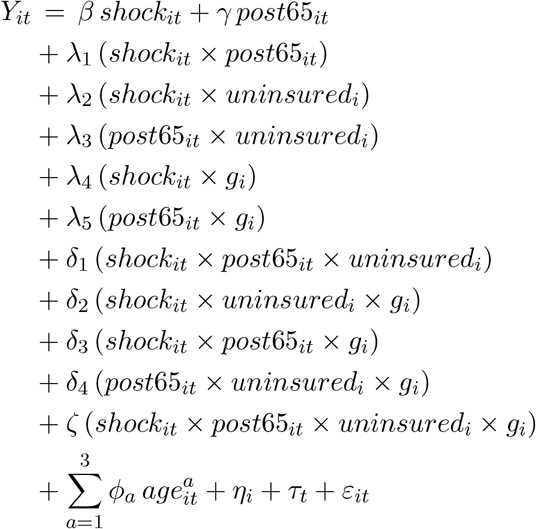

**Effect of the health shock on the outcome for the previously uninsured** To calculate the effect of the shock on the outcome, we evaluate the derivative of the outcome with respect to shock:

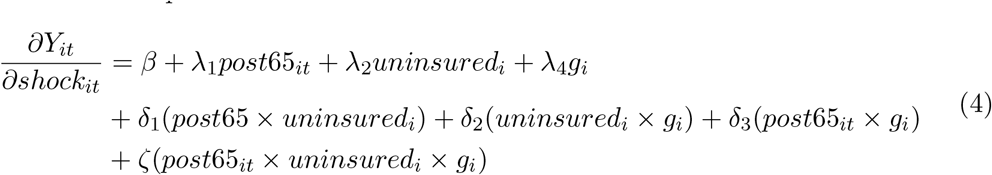

We can look at the decomposition for the different genotypes (high and low PGS) and shock timing (before and after 65):

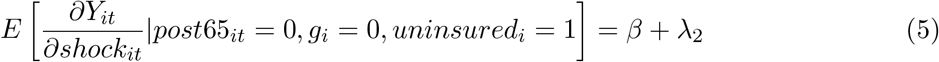

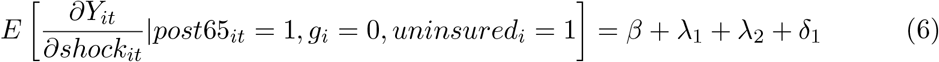

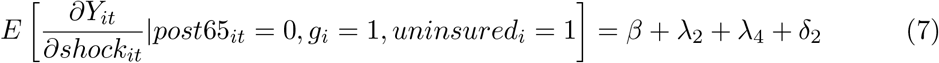

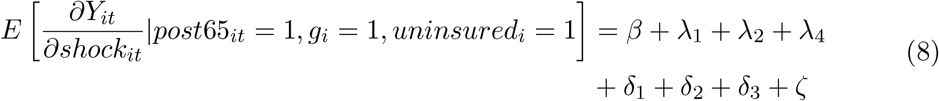

Calculating the first two differences as above:

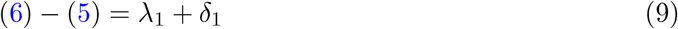

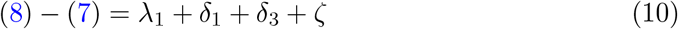

And finally the genetic heterogeneity in this difference:

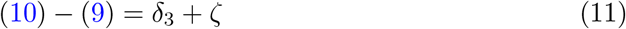

## C.5.2 OLS estimation: Low, middle and high PGS

Current smoking status (*Y*) is regressed on the full set of interactions between the indicators for the health shock (*shock*), being uninsured pre-65 (*uninsured*), Medicare eligibility (*post*65), medium polygenic risk for smoking (*g^m^*) and high polygenic risk for smoking (*g^h^*):

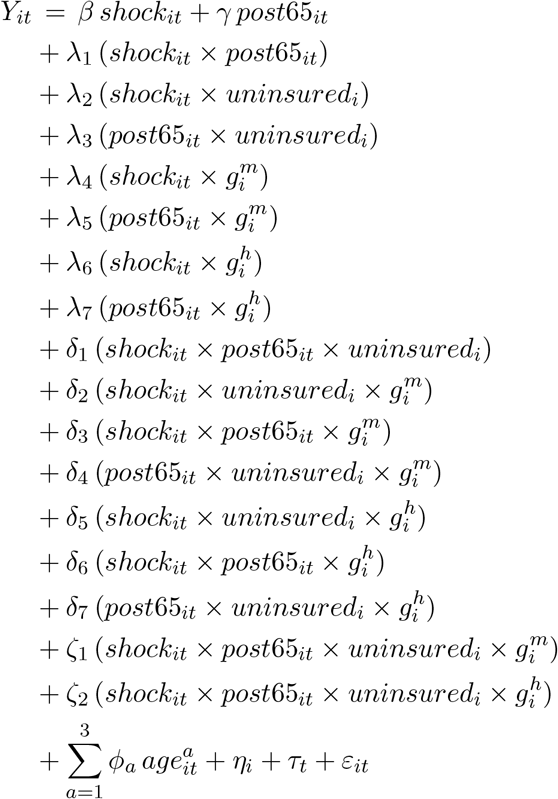

**Effect of the shock on the outcome** The derivative of the outcome with respect to shock is:

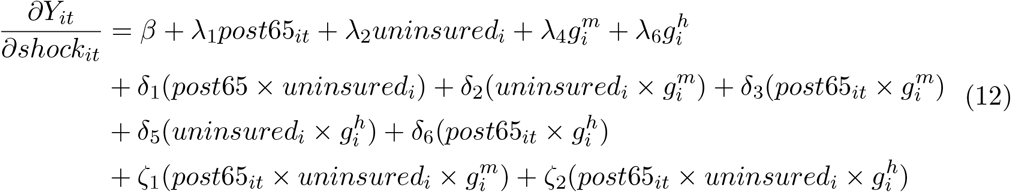

Again, we can look at the decomposition:

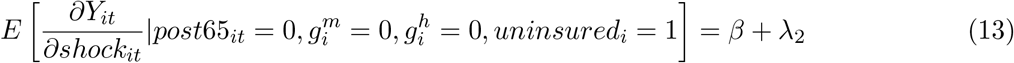

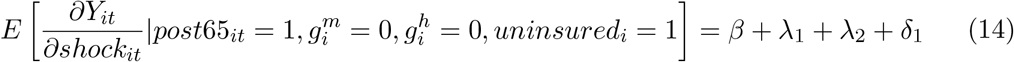

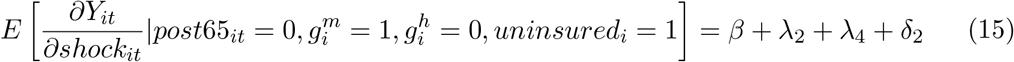

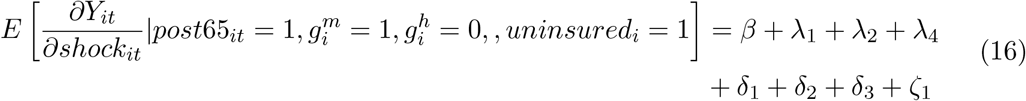

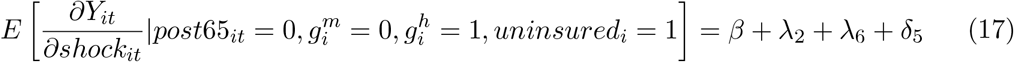

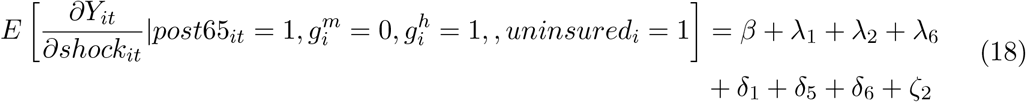

Calculating the first two differences as above::

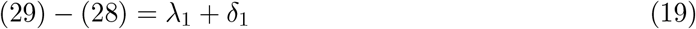

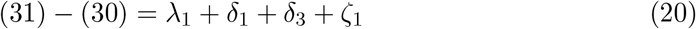

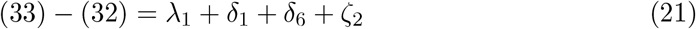

And again the genetic heterogeneity in this difference:

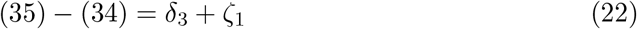

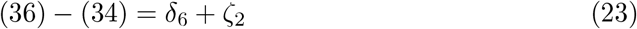

## D Results

## D.1 Sample Characteristics

Table 5 displays summary statistics for the subset of the study sample that experienced a cardiovascular health shock over the course of the observation period, stratified by timing of the shock (pre-65 versus post-65) and genetic group. Within a genetic group, demographics are mostly similar across the timing strata. However, in both groups, those experiencing the shock after age 65 are on average older at baseline. In the high-PGS group, there are also relatively more women experiencing the shock after the age of 65 than before 65. This is consistent with the general pattern that women experience cardiovascular disease later in life than men (Lloyd-Jones et al., 2010).

The subset of respondents affected by cardiovascular illness during the observed years differed in some characteristics from those unaffected. Table 6 shows a comparison.

## D.2 Main Results

Table 7 reports the covariance matrix of these estimated coefficients. These coefficients and standard errors are used to calculate the effect of health shocks on the smoking probability of individuals who are uninsured before the age of 65—the subgroup of interest—for the 4 combinations of shock timing (before or after 65) and polygenic score (high or low). The derivation of these effects is described in Appendix Section C.5.

**Table D.5:**
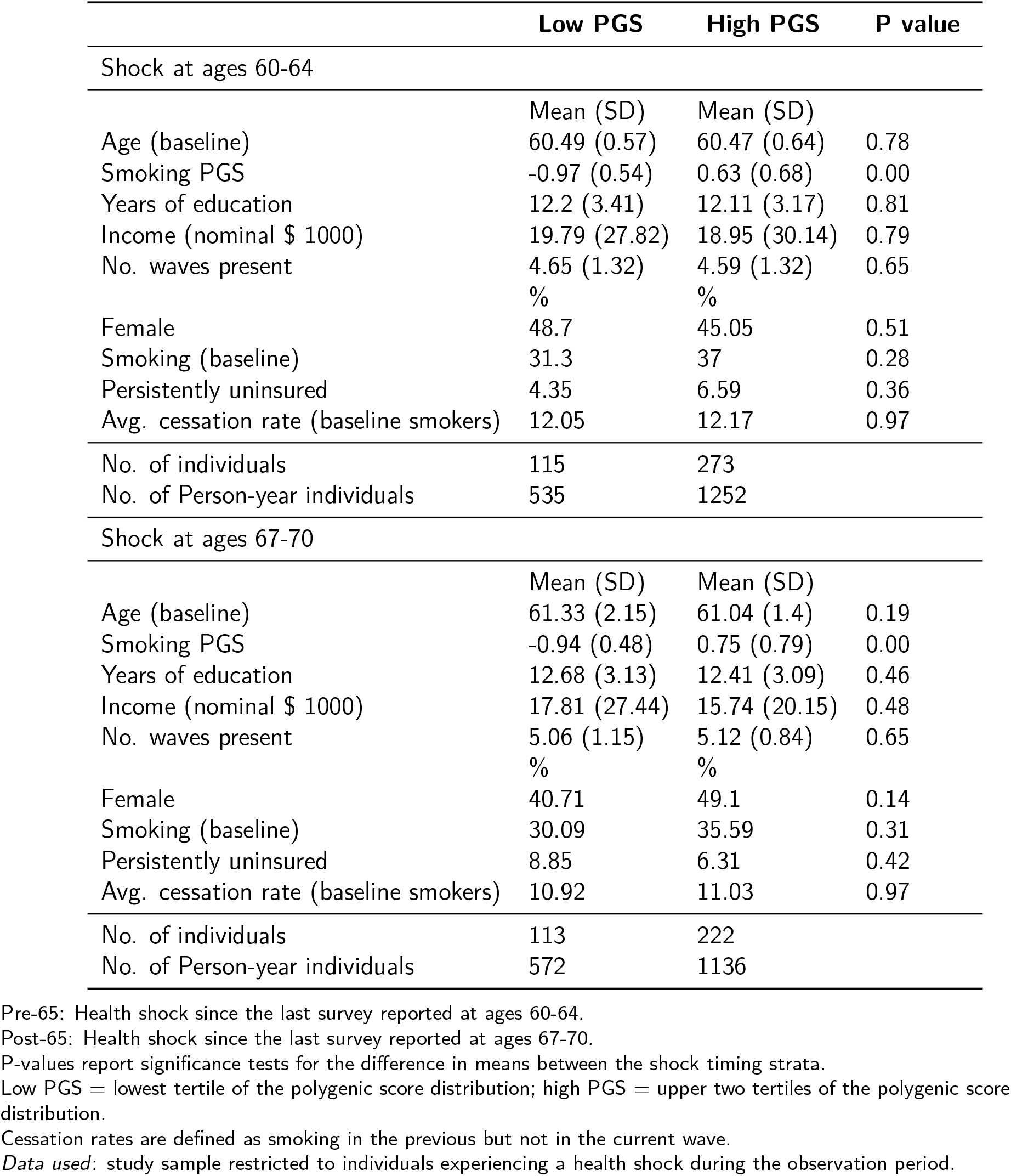
Descriptive Statistics for the Subset of the study sample with a Health Shock, Stratified by Timing of the Shock and Genetic Group

|  | Low PGS | High PGS | P value |
| --- | --- | --- | --- |
| Shock at ages 60-64 |  |  |  |
|  | Mean (SD) | Mean (SD) |  |
| Age (baseline) | 60.49 (0.57) | 60.47 (0.64) | 0.78 |
| Smoking PGS | -0.97 (0.54) | 0.63 (0.68) | 0.00 |
| Years of education | 12.2 (3.41) | 12.11 (3.17) | 0.81 |
| Income (nominal \$ 1000) | 19.79 (27.82) | 18.95 (30.14) | 0.79 |
| No. waves present | 4.65 (1.32) | 4.59 (1.32) | 0.65 |
|  | % | % |  |
| Female | 48.7 | 45.05 | 0.51 |
| Smoking (baseline) | 31.3 | 37 | 0.28 |
| Persistently uninsured | 4.35 | 6.59 | 0.36 |
| Avg. cessation rate (baseline smokers) | 12.05 | 12.17 | 0.97 |
| No. of individuals | 115 | 273 |  |
| No. of Person-year individuals | 535 | 1252 |  |
| Shock at ages 67-70 |  |  |  |
|  | Mean (SD) | Mean (SD) |  |
| Age (baseline) | 61.33 (2.15) | 61.04 (1.4) | 0.19 |
| Smoking PGS | -0.94 (0.48) | 0.75 (0.79) | 0.00 |
| Years of education | 12.68 (3.13) | 12.41 (3.09) | 0.46 |
| Income (nominal \$ 1000) | 17.81 (27.44) | 15.74 (20.15) | 0.48 |
| No. waves present | 5.06 (1.15) | 5.12 (0.84) | 0.65 |
|  | % | % |  |
| Female | 40.71 | 49.1 | 0.14 |
| Smoking (baseline) | 30.09 | 35.59 | 0.31 |
| Persistently uninsured | 8.85 | 6.31 | 0.42 |
| Avg. cessation rate (baseline smokers) | 10.92 | 11.03 | 0.97 |
| No. of individuals | 113 | 222 |  |
| No. of Person-year individuals | 572 | 1136 |  |

Pre-65: Health shock since the last survey reported at ages 60-64.
Post-65: Health shock since the last survey reported at ages 67-70.
P-values report significance tests for the difference in means between the shock timing strata.
Low PGS = lowest tertile of the polygenic score distribution; high PGS = upper two tertiles of the polygenic score distribution.
Cessation rates are defined as smoking in the previous but not in the current wave.
Data used: study sample restricted to individuals experiencing a health shock during the observation period.

**Table D.6:** Descriptive Statistics for the Study Sample, Stratified by Future Health Shock Status

|  | All | New Shock | No new Shock | P value |
| --- | --- | --- | --- | --- |
|  | Mean (SD) | Mean (SD) | Mean (SD) |  |
| Age (baseline) | 61.15 (1.87) | 60.74 (1.14) | 61.21 (1.94) | 0.00 |
| Smoking PGS | 0.1 (0.99) | 0.18 (1.02) | 0.09 (0.99) | 0.03 |
| Years of education | 12.48 (3.09) | 12.33 (3.16) | 12.5 (3.08) | 0.18 |
| Income (nominal \$ 1000) | 20.41 (34.97) | 17.97 (26.62) | 20.75 (35.98) | 0.01 |
| No. waves present | 4.45 (1.36) | 4.85 (1.15) | 4.39 (1.38) | 0.00 |
|  | % | % | % |  |
| Female | 49.86 | 45.9 | 50.42 | 0.02 |
| Smoking (baseline) | 29.5 | 34.08 | 28.86 | 0.01 |
| Persistently uninsured | 5.88 | 6.4 | 5.8 | 0.54 |
| CV health shock | 12.28 | 100 | 0 | - |
| Avg. cessation rate (baseline smokers) | 10.3 | 11.35 | 10.11 | 0.21 |
| No. of individuals | 5854 | 719 | 5135 |  |

Health shock: diagnosed with a new cardiovascular condition during the time since the last HRS survey, but no history of cardiovascular disease prior to this diagnosis. P-values report significance tests for the difference in means between the health shock strata. Cessation rates were defined as smoking in the previous but not in the current wave.
*Data used:* HRS study sample, $n = 5,854$ .

## D.3 Robustness Checks

## D.3.1 Using all tertiles

In this section, we derive the results using all of the three tertiles of the PGS distribution, as derived in Appendix Section D.3.1. Results shown in Tables 8 and Figure 12, show that the effects for the upper two tertiles of the polygenic score are virtually the same. To improve statistical power and simplicity of exposition, the main results are always presented by pooling these two tertiles toghether into a single category labeled high-PGS.

## D.3.2 Median Split of the polygenic score

In this section, we derive the results by using a median-split for the PGS, instead of tertiles. Results shown in Tables 9 10, as well as Figure 13, show that the effects are virtually the same, albeit a bit smaller in magnitude and less precisely estimated, as when splitting the PGS according to tertiles.

## D.3.3 Older GWAS Summary Statistics

In this section, we also use the polygenic score publicly provided by the HRS (Ware, Schmitz and Faul, 2017) for the smoking phenotype “regular smoking” (having smoked more than 100 cigarettes throughout one’s life). This score is constructed as a weighted sum of the genotype over the 779,538 SNPs that overlap between the HRS genetic database and a 2010 GWAS meta-analysis conducted by the Tobacco and Genetics Consortium (The Tobacco and Genetics Consortium et al., 2010).

**Figure D.12:**
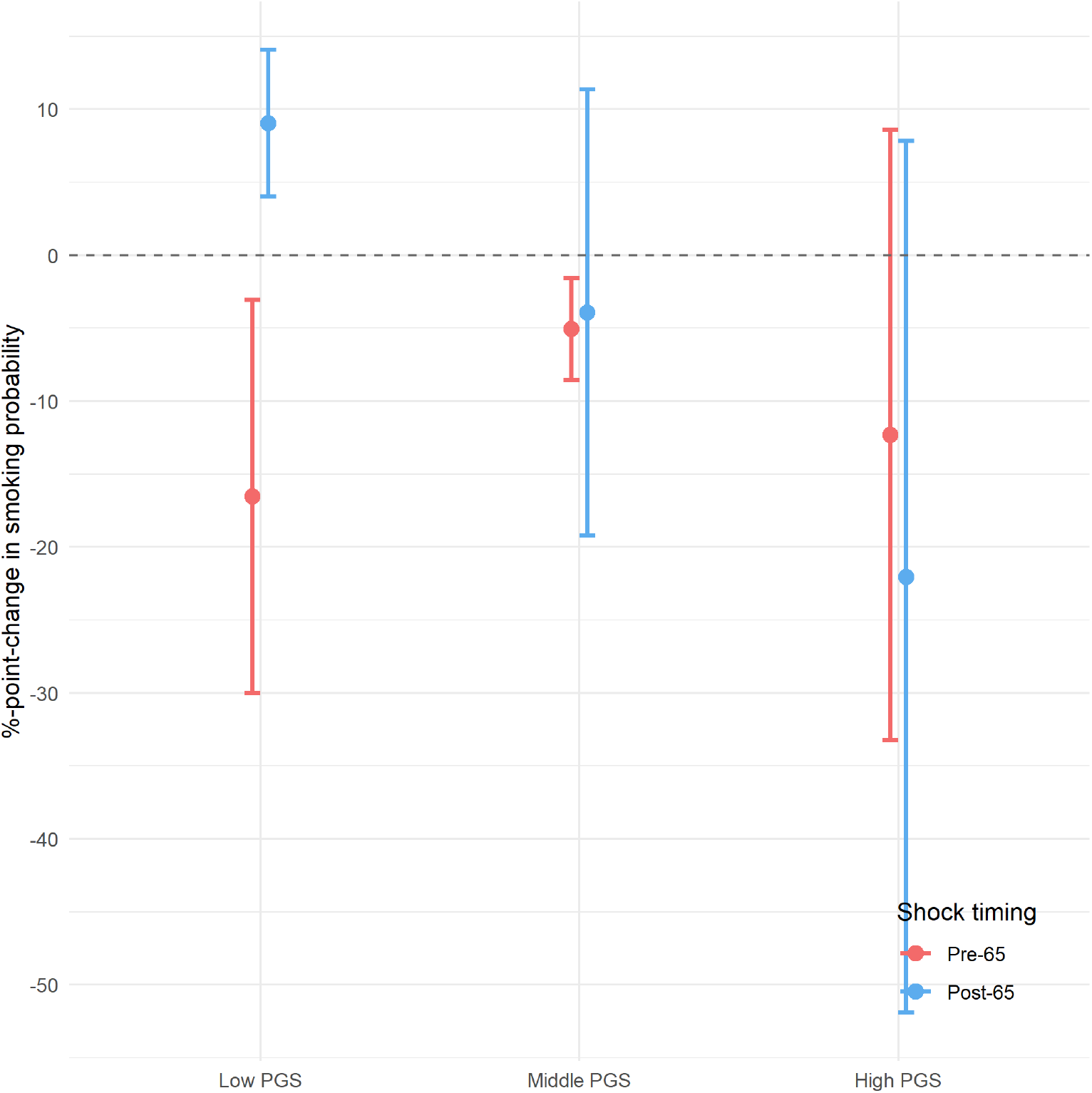
Effect of a Health Shock on the Smoking Probability in the Pre-65 Uninsured Subgroup, Stratified by Timing of the Shock and Genetic Type (median split) *Notes:* Low PGS = lowest tertile of the polygenic score distribution; Middle PGS = high PGS = upper two tertiles of the polygenic score distribution. Pre-65: Health shock since the last survey reported at ages 60-64. Post-65: Health shock since the last survey reported at ages 67-70. Estimates and standard errors are shown in Panel A of Table 2. Effects are estimated using the coefficients in Table 3 and following the derivation described in C.5. Bars show 95% confidence intervals, standard error clustered at the individual level. *Data used*: HRS study sample, n = 5,854. Supplement.

Results shown in Tables 11 and Figure 14, show that the effects are virtually the same, possibly even a bit stronger.

## D.3.4 Using different polygenic scores

Other polygenic scores (PGS), besides the one for being a smoker, might be driving this heterogeneity in moral hazard. We estimated the same analysis outlined in equation 2 but replacing *g_i_* with the following PGS: the PGS for cigarettes per day (CPD) (Liu et al., 2019), the PGS for educational attainment and the one for cognitive abilities (Lee et al., 2018), the PGS for risk tolerance (Karlsson Linnér et al., 2019), the PGS for non-cognitive skills (Demange et al., 2020), and the PGS for Body-Mass-Index (Yengo et al., 2018). We chose polygenic scores for traits that are genetically correlated with smoking, such as CPD and risk tolerance, or plausibly related to strategic behaviors and moral hazard, such as education, cognition, and non-cognitive skills. BMI is meant more as a placebo.

**Figure D.13:**
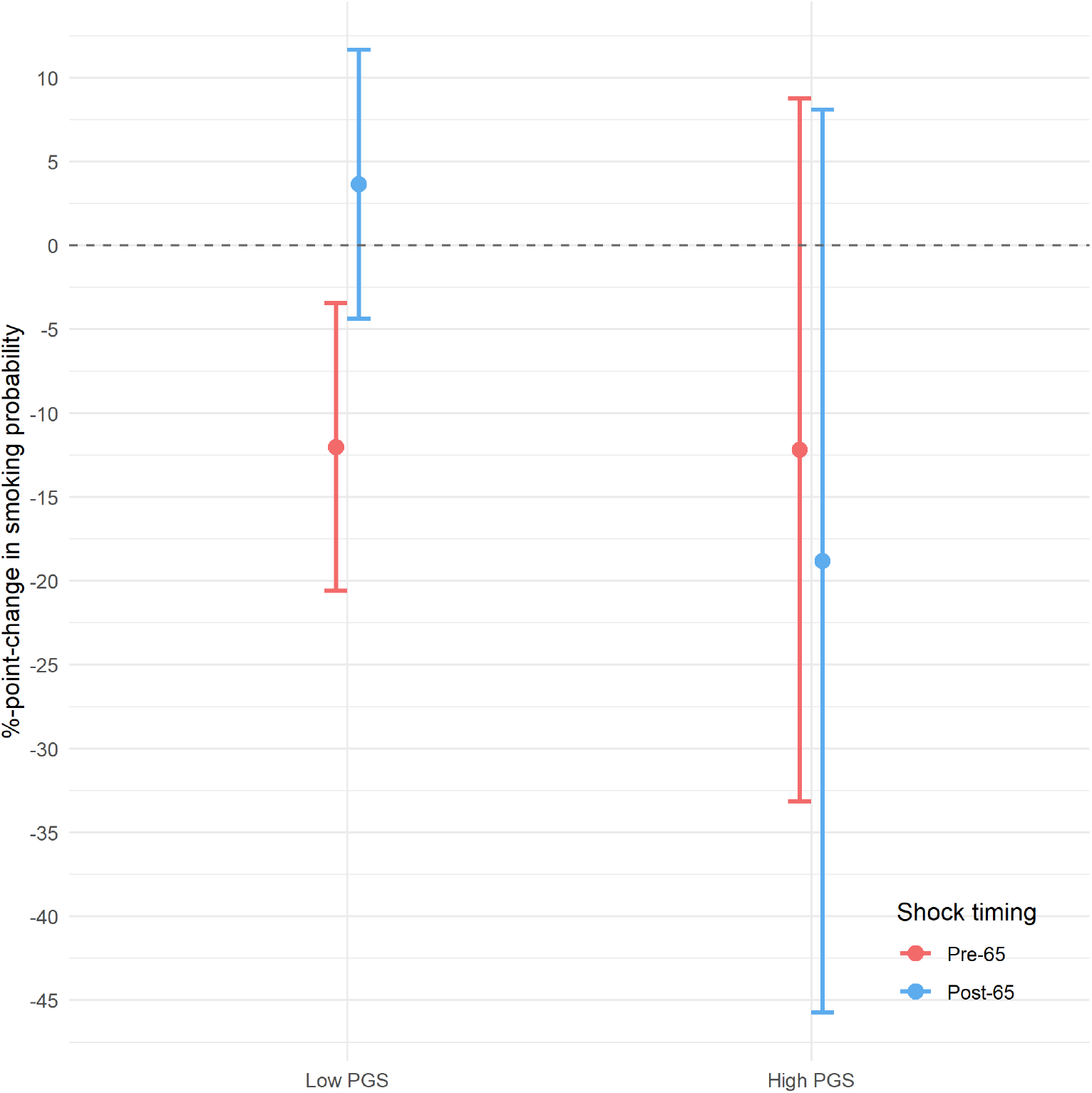
Effect of a Health Shock on the Smoking Probability in the Pre-65 Uninsured Subgroup, Stratified by Timing of the Shock and Genetic Type (median split) *Notes:* Low PGS = polygenic score below the median; high PGS = polygenic score above the median. Pre-65: Health shock since the last survey reported at ages 60-64. Post-65: Health shock since the last survey reported at ages 67-70. Estimates and standard errors are shown in Panel A of Table 2. Effects are estimated using the coefficients in Table 3 and following the derivation described in C.5. Bars show 95% confidence intervals, standard error clustered at the individual level. *Data used*: HRS study sample, n = 5,854.

**Figure D.14:**
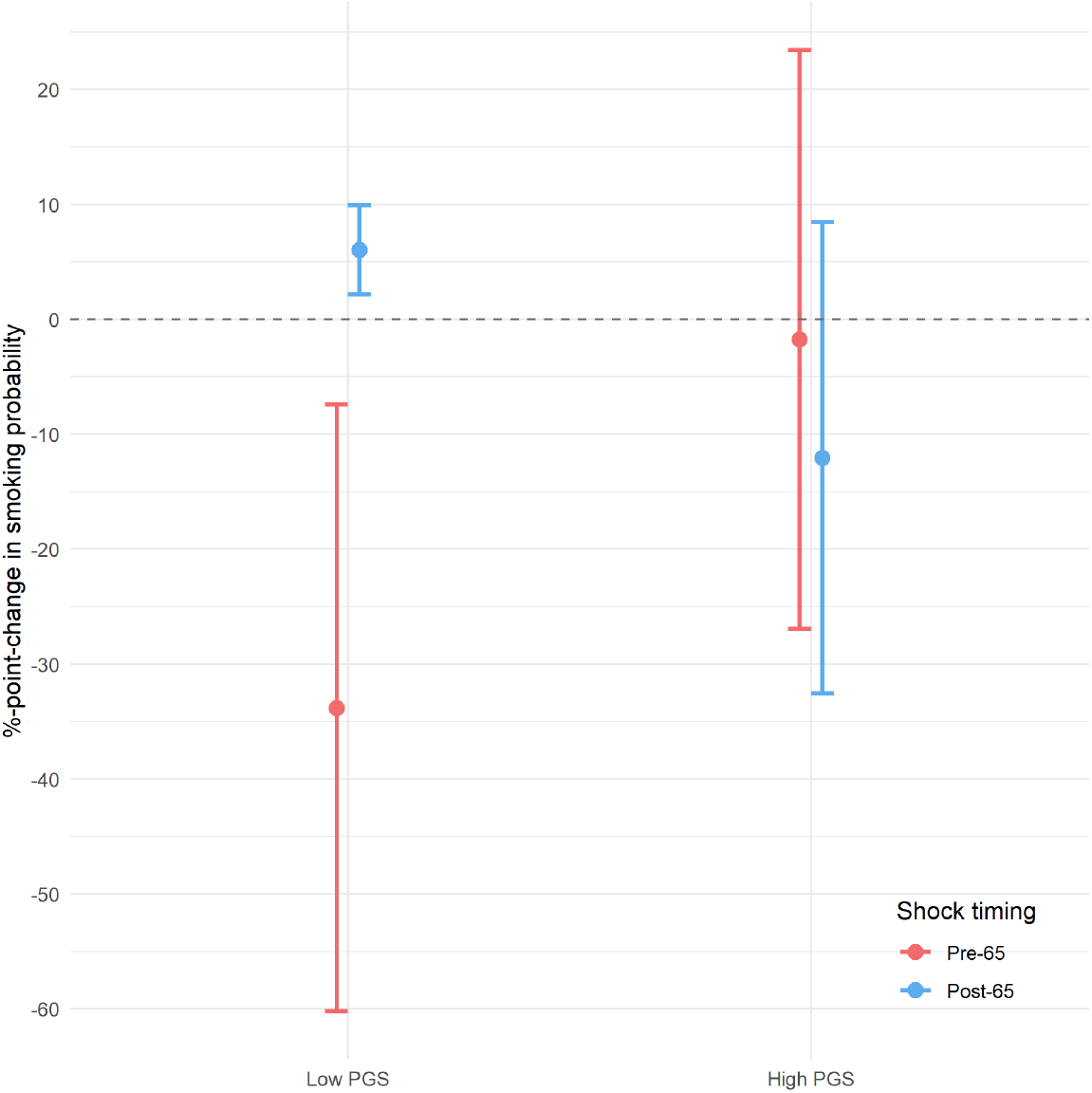
Effect of a Health Shock on the Smoking Probability in the Pre-65 Uninsured Subgroup, Stratified by Timing of the Shock and Genetic Type (older PGS) *Notes:* Low PGS = polygenic score below the median; high PGS = polygenic score above the median. Pre-65: Health shock since the last survey reported at ages 60-64. Post-65: Health shock since the last survey reported at ages 67-70. Estimates and standard errors are shown in Panel A of Table 2. Effects are estimated using the coefficients in Table 3 and following the derivation described in C.5. Bars show 95% confidence intervals, standard error clustered at the individual level. *Data used*: HRS study sample, n = 5,854.

**Table D.7:**
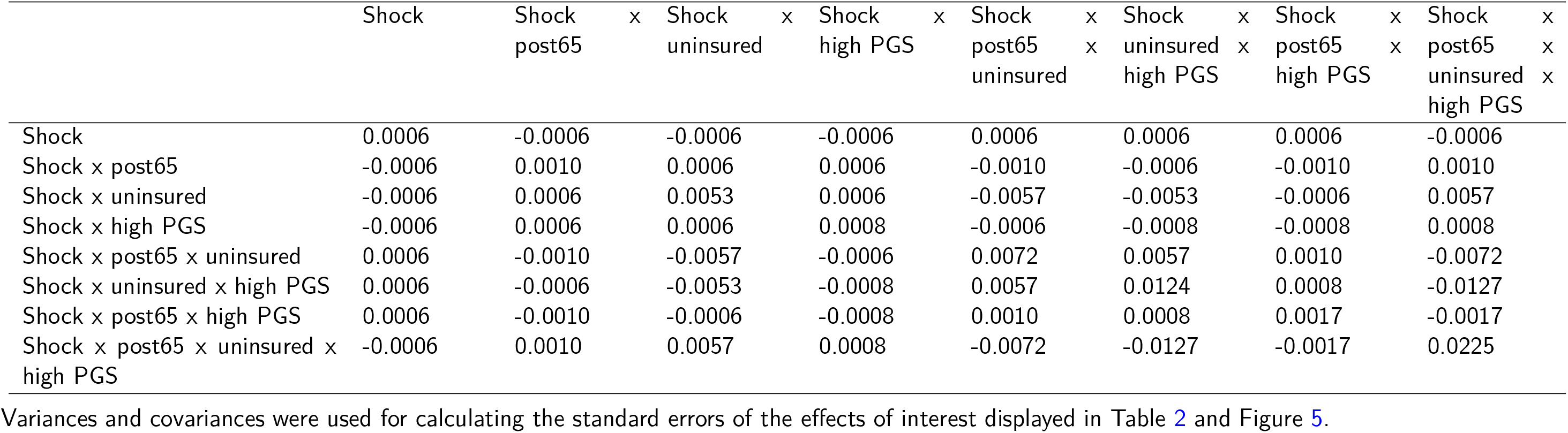
Covariance Matrix for Regression Coefficients in the last column of Table 3

**Table D.8:** Summary of Statistical Results for the Pre-65 Uninsured Subgroup, Stratified by Timing of the Shock and Genetic Group (Median)

| **Effect of health shock on smoking probability** |  |  |  |
| --- | --- | --- | --- |
|  | Low PGS | Middle PGS | High PGS |
| Pre 65 | -0.165**<br>(0.069) | -0.051***<br>(0.018) | -0.123<br>(0.107) |
| Post 65 | 0.09*** | -0.039 | -0.221 |
| **Effect of health insurance on effect of health shock** |  |  |  |
|  | (0.026) | (0.078) | (0.152) |
|  | Low PGS | Middle PGS | High PGS |
| Post 65 - Pre 65 | 0.256*** | 0.011 | -0.097 |
| **Differential effect of health insurance by genetic group** |  |  |  |
|  | (0.079) | (0.076) | (0.185) |
|  | High PGS - low PGS | Middle PGS - low PGS | High PGS - middle PGS |
| Post 65 - Pre 65 | -0.353*<br>(0.201) | -0.245**<br>(0.109) | -0.109<br>(0.2) |

*Data used:* HRS study sample, $n = 5,854$ .

**Table D.9:**
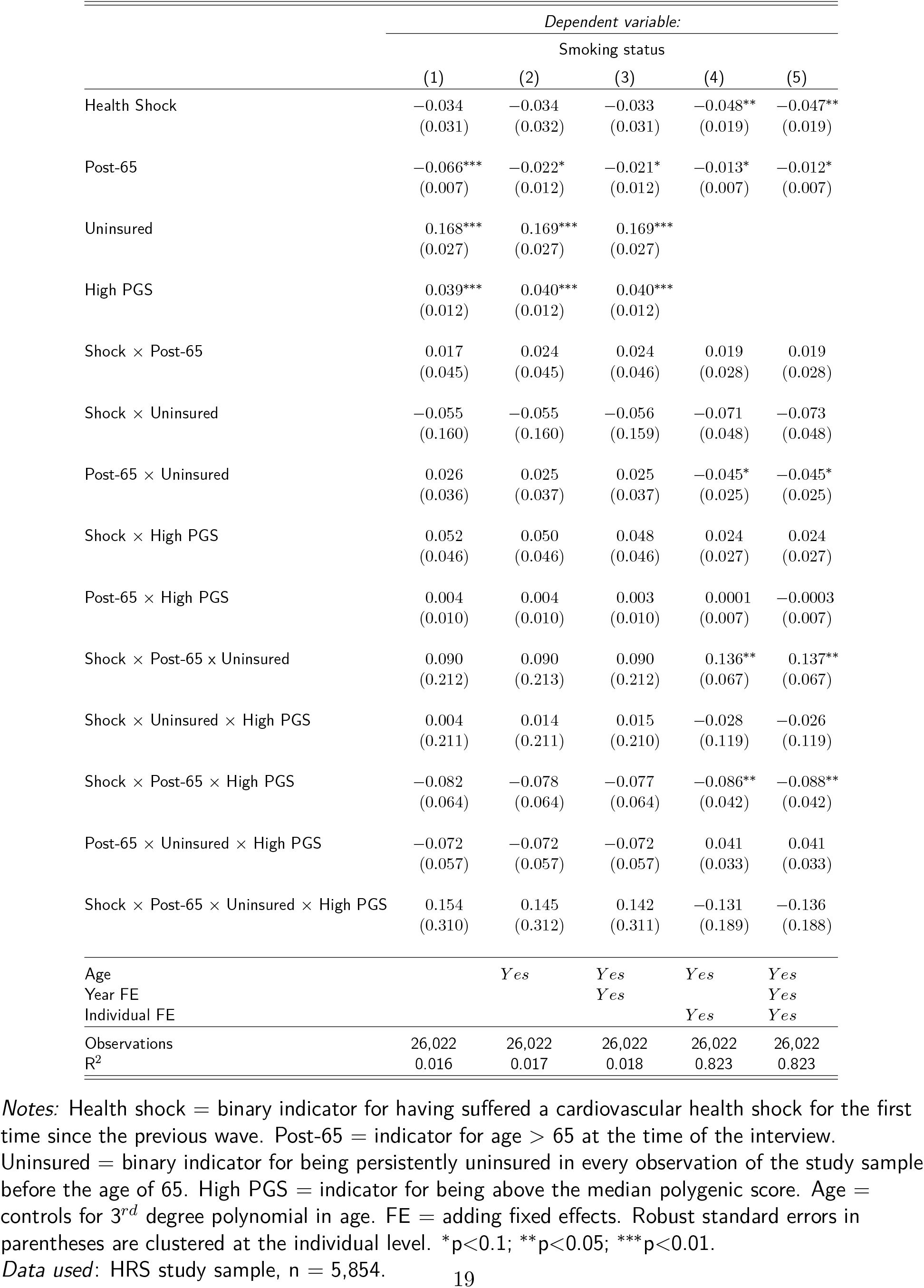
Coefficients from estimating the linear probability model in equation (2) using OLS (PGS median split)

|  | *Dependent variable:* |  |  |  |  |
| --- | --- | --- | --- | --- | --- |
|  | Smoking status |  |  |  |  |
|  | (1) | (2) | (3) | (4) | (5) |
| Health Shock | −0.034<br>(0.031) | −0.034<br>(0.032) | −0.033<br>(0.031) | −0.048**<br>(0.019) | −0.047**<br>(0.019) |
| Post-65 | −0.066***<br>(0.007) | −0.022*<br>(0.012) | −0.021*<br>(0.012) | −0.013*<br>(0.007) | −0.012*<br>(0.007) |
| Uninsured | 0.168***<br>(0.027) | 0.169***<br>(0.027) | 0.169***<br>(0.027) |  |  |
| High PGS | 0.039***<br>(0.012) | 0.040***<br>(0.012) | 0.040***<br>(0.012) |  |  |
| Shock × Post-65 | 0.017<br>(0.045) | 0.024<br>(0.045) | 0.024<br>(0.046) | 0.019<br>(0.028) | 0.019<br>(0.028) |
| Shock × Uninsured | −0.055<br>(0.160) | −0.055<br>(0.160) | −0.056<br>(0.159) | −0.071<br>(0.048) | −0.073<br>(0.048) |
| Post-65 × Uninsured | 0.026<br>(0.036) | 0.025<br>(0.037) | 0.025<br>(0.037) | −0.045*<br>(0.025) | −0.045*<br>(0.025) |
| Shock × High PGS | 0.052<br>(0.046) | 0.050<br>(0.046) | 0.048<br>(0.046) | 0.024<br>(0.027) | 0.024<br>(0.027) |
| Post-65 × High PGS | 0.004<br>(0.010) | 0.004<br>(0.010) | 0.003<br>(0.010) | 0.0001<br>(0.007) | −0.0003<br>(0.007) |
| Shock × Post-65 × Uninsured | 0.090<br>(0.212) | 0.090<br>(0.213) | 0.090<br>(0.212) | 0.136**<br>(0.067) | 0.137**<br>(0.067) |
| Shock × Uninsured × High PGS | 0.004<br>(0.211) | 0.014<br>(0.211) | 0.015<br>(0.210) | −0.028<br>(0.119) | −0.026<br>(0.119) |
| Shock × Post-65 × High PGS | −0.082<br>(0.064) | −0.078<br>(0.064) | −0.077<br>(0.064) | −0.086**<br>(0.042) | −0.088**<br>(0.042) |
| Post-65 × Uninsured × High PGS | −0.072<br>(0.057) | −0.072<br>(0.057) | −0.072<br>(0.057) | 0.041<br>(0.033) | 0.041<br>(0.033) |
| Shock × Post-65 × Uninsured × High PGS | 0.154<br>(0.310) | 0.145<br>(0.312) | 0.142<br>(0.311) | −0.131<br>(0.189) | −0.136<br>(0.188) |
| Age |  | *Yes* | *Yes* | *Yes* | *Yes* |
| Year FE |  |  | *Yes* |  | *Yes* |
| Individual FE |  |  |  | *Yes* | *Yes* |
| Observations | 26,022 | 26,022 | 26,022 | 26,022 | 26,022 |
| R<sup>2</sup> | 0.016 | 0.017 | 0.018 | 0.823 | 0.823 |

*Notes:* Health shock = binary indicator for having suffered a cardiovascular health shock for the first time since the previous wave. Post-65 = indicator for age > 65 at the time of the interview. Uninsured = binary indicator for being persistently uninsured in every observation of the study sample before the age of 65. High PGS = indicator for being above the median polygenic score. Age = controls for 3<sup>rd</sup> degree polynomial in age. FE = adding fixed effects. Robust standard errors in parentheses are clustered at the individual level. \*p<0.1; \*\*p<0.05; \*\*\*p<0.01.
*Data used:* HRS study sample, n = 5,854.

**Table D.10:** Summary of Statistical Results for the Pre-65 Uninsured Subgroup, Stratified by Timing of the Shock and Genetic Group (Median)

| **Effect of health shock on smoking probability** |  |  |
| --- | --- | --- |
|  | Low PGS | High PGS |
| Pre 65 | -0.12***<br>(0.044) | -0.122<br>(0.107) |
| Post 65 | 0.036<br>(0.041) | -0.188<br>(0.137) |
| **Effect of health insurance on effect of health shock** |  |  |
|  | Low PGS | High PGS |
| Post 65 - Pre 65 | 0.157**<br>(0.061) | -0.066<br>(0.173) |
| **Differential effect of health insurance by genetic group** |  |  |
|  | High PGS - low PGS |  |
| Post 65 - Pre 65 | -0.223<br>(0.184) |  |

*Data used:* HRS study sample, n = 5,854.

**Table D.11:** Summary of Statistical Results for the Pre-65 Uninsured Subgroup, Stratified by Timing of the Shock and Genetic Group (older PGS)

| **Effect of health shock on smoking probability** |  |  |
| --- | --- | --- |
|  | Low PGS | High PGS |
| Pre 65 | -0.338**<br>(0.135) | -0.018<br>(0.128) |
| Post 65 | 0.06***<br>(0.02) | -0.12<br>(0.105) |
| **Effect of health insurance on effect of health shock** |  |  |
|  | Low PGS | High PGS |
| Post 65 - Pre 65 | 0.399***<br>(0.139) | -0.103<br>(0.163) |
| **Differential effect of health insurance by genetic group** |  |  |
|  | High PGS - low PGS |  |
| Post 65 - Pre 65 | -0.501**<br>(0.214) |  |

*Data used:* HRS study sample, n = 5,854.
*Data used:* HRS study sample, $n = 5,854$ .

**Figure D.15:**
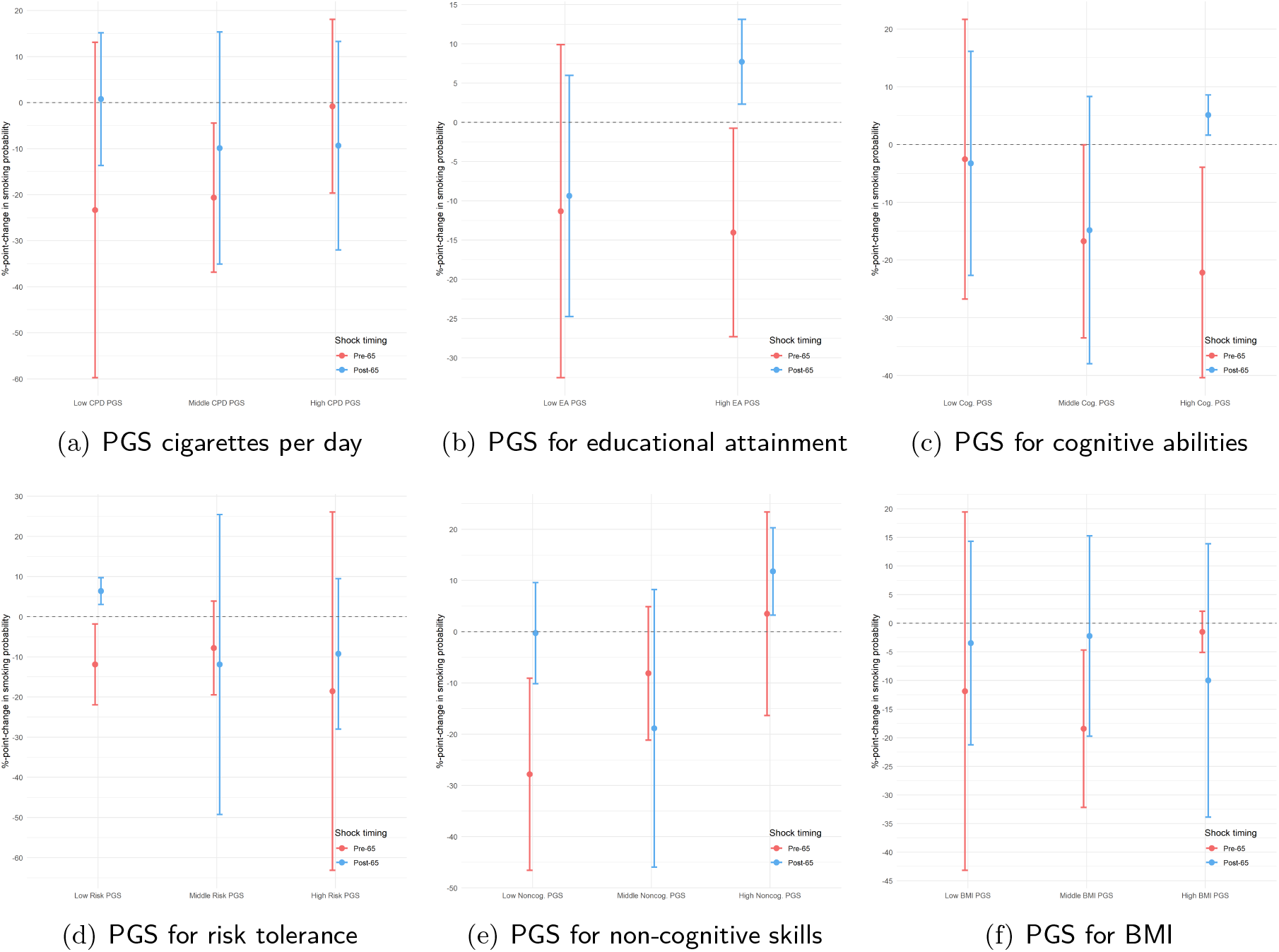
Other Polygenic Scores as Proxy for Moral Hazard Heterogeneity *Notes:* The figures report the effect of suffering a health shock on the probability of smoking for the pre-65 uninsured subgroup, stratified by timing of the shock (before and after the age of 65) and different polygenic scores. Pre-65: Health shock since the last survey reported at ages 60-64. Post-65: Health shock since the last survey reported at ages 67-70. Effects are estimated using a combination of the coefficients from equation 2 where *g_i_* is replaced by the different polygenic scores reported in the sub-figure captions, following the derivation described in C.5. Bars show 95% confidence intervals, standard error clustered at the individual level. *Data used*: HRS study sample, n = 5,854.

## D.3.5 Relaxing the Criteria for Inclusion in the Pre-65 Uninsured Group

**Table D.12:** Summary of Statistical Results for the Pre-65 Uninsured Subgroup (Using Different Definitions of the Pre-65 Uninsured Status Indicator)

|  | 100% uninsured |  | 66% uninsured |  | 33% uninsured |  |
| --- | --- | --- | --- | --- | --- | --- |
| Effect of health shock on smoking probability |  |  |  |  |  |  |
|  | Low PGS | High PGS | Low PGS | High PGS | Low PGS | High PGS |
| Pre 65 | -0.165**<br>(0.069) | -0.108<br>(0.083) | -0.147**<br>(0.059) | -0.055<br>(0.06) | -0.055<br>(0.068) | -0.09*<br>(0.048) |
| Post 65 | 0.09***<br>(0.026) | -0.13<br>(0.089) | 0.086***<br>(0.021) | -0.091<br>(0.078) | 0.052<br>(0.036) | -0.102**<br>(0.05) |
| Effect of health insurance on effect of health shock |  |  |  |  |  |  |
|  | Low PGS | High PGS | Low PGS | High PGS | Low PGS | High PGS |
| Post 65 - Pre 65 | 0.256***<br>(0.079) | -0.023<br>(0.121) | 0.233***<br>(0.067) | -0.036<br>(0.098) | 0.107<br>(0.076) | -0.012<br>(0.069) |
| Differential effect of health insurance by genetic group |  |  |  |  |  |  |
|  | High PGS | - low PGS | High PGS | - low PGS | High PGS | - low PGS |
| Post 65 - Pre 65 | -0.279*<br>(0.144) |  | -0.269**<br>(0.118) |  | -0.119<br>(0.103) |  |

Main study results, for comparison.
Pre-65 uninsured status indicator set to 1 for respondents who were uninsured in at least 66% of all pre-65 observations.
Pre-65 uninsured status indicator set to 1 for respondents who were uninsured in at least 33% of all pre-65 observations.
\*P < 0.1; \*\*P < 0.05; \*\*\*P < 0.01. Robust standard errors in parentheses were clustered at the individual level.
Low PGS = lowest tertile of the polygenic score distribution; high PGS = upper two tertiles of the polygenic score distribution.
Pre-65: Health shock since the last survey reported at ages 60-64. Post-65: Health shock since the last survey reported at ages 67-70.
Data used: HRS study sample, n = 5,854.

## D.3.6 Including Individuals with Shocks Reported at Ages 65 and 66

**Table D.13:** Summary of Statistical Results for the Pre-65 Uninsured Subgroup (Including Individuals Reporting a Health Shock when Aged 65 or 66 in the study Sample)

|  | *Analytic Sample* |  | *Including reported at 65/66* |  |
| --- | --- | --- | --- | --- |
| Effect of health shock on smoking probability |  |  |  |  |
|  | Low PGS | High PGS | Low PGS | High PGS |
| Pre 65 | -0.165**<br>(0.069) | -0.108<br>(0.083) | -0.166**<br>(0.069) | -0.113<br>(0.084) |
| Post 65 | 0.09***<br>(0.026) | -0.13<br>(0.089) | 0.091***<br>(0.025) | -0.213**<br>(0.089) |
| Effect of health insurance on effect of health shock |  |  |  |  |
|  | Low PGS | High PGS | Low PGS | High PGS |
| Post 65 - Pre 65 | 0.256***<br>(0.079) | -0.023<br>(0.121) | 0.257***<br>(0.079) | -0.1<br>(0.121) |
| Differential effect of health insurance by genetic group |  |  |  |  |
|  | High PGS | - low PGS | High PGS | - low PGS |
| Post 65 - Pre 65 | -0.279*<br>(0.144) |  | -0.356**<br>(0.145) |  |

Main study results, for comparison.
study sample additionally includes individuals who reported experiencing a health shock since the last survey wave when interviewed at ages 65 or 66.
\*P < 0.1; \*\*P < 0.05; \*\*\*P < 0.01. Robust standard errors in parentheses were clustered at the individual level.
Low PGS = lowest tertile of the polygenic score distribution; high PGS = upper two tertiles of the polygenic score distribution.
Pre-65: Health shock since the last survey reported at ages 60-64. Post-65: Health shock since the last survey reported at ages 67-70.
*Data used:* study sample, but skipping Step 6 in Section C.2.1.

## D.3.7 Using Medicare Enrollment Status instead of age 65

Medicare enrollment refers to the RAND variable GOVMR, which indicates whether the respondent is covered by Medicare in a given wave. For details on the survey questions and construction of this variable, see the RAND HRS documentation.

**Table D.14:** Summary of Statistical Results for the Pre-65 Uninsured Subgroup (Using Medicare Enrollment Status Instead of Medicare Eligibility Status)

| **Effect of health shock on smoking probability** |  |  |
| --- | --- | --- |
|  | Low PGS | High PGS |
| Without Medicare | -0.136**<br>(0.066) | -0.109<br>(0.083) |
| With Medicare | 0.097***<br>(0.028) | -0.124<br>(0.089) |
| **Effect of health insurance on effect of health shock** |  |  |
|  | Low PGS | High PGS |
| With - Without Medicare | 0.233***<br>(0.077) | -0.016<br>(0.121) |
| **Differential effect of health insurance by genetic group** |  |  |
|  | High PGS | - low PGS |
| With - Without Medicare | -0.248*<br>(0.144) |  |

\* $P < 0.1$ ; \*\* $P < 0.05$ ; \*\*\* $P < 0.01$ . Robust standard errors in parentheses, clustered at the individual level. Low PGS = lowest tertile of the polygenic score distribution; high PGS = upper two tertiles of the polygenic score distribution. Effects are estimated using the coefficients in Table 3 and following the derivation described in C.5, but replacing the post-65 indicator with Medicare enrollment status. *Data used*: HRS study sample with the additional restriction of a non-missing Medicare enrollment status.

## D.4 Confounders

Tables 15–17 report the results for the effect of the shock on potential confounding variables.

## E The Model

In this section, we introduce a model to explore the theoretical basis and the testable empirical consequences of heterogeneity in moral hazard.^16^

What drives heterogeneity in moral hazard? In the context of health insurance, moral hazard is usually considered to be a reaction to insurance coverage: an increase in insurance coverage leads to changes in health behaviors, such as increased usage of medical care (Einav and Finkelstein, 2018) or smoking. Hence moral hazard can be considered as a sort of price sensitivity of the agent (as in Einav et al., 2013). Knowing they can afford to go to the doctor if they fall sick, the insured agents might be more inclined to engage in immediately rewarding behavior which is harmful in the long run, such as smoking.^17^ Heterogeneity in this response to health care coverage can then be driven by social factors, like exposure to family or peers who behave similarly (Chatterjee et al., 2018; Hoffmann, 2017), or biological factors, like genetic propensity to smoke.

**Table D.15:**
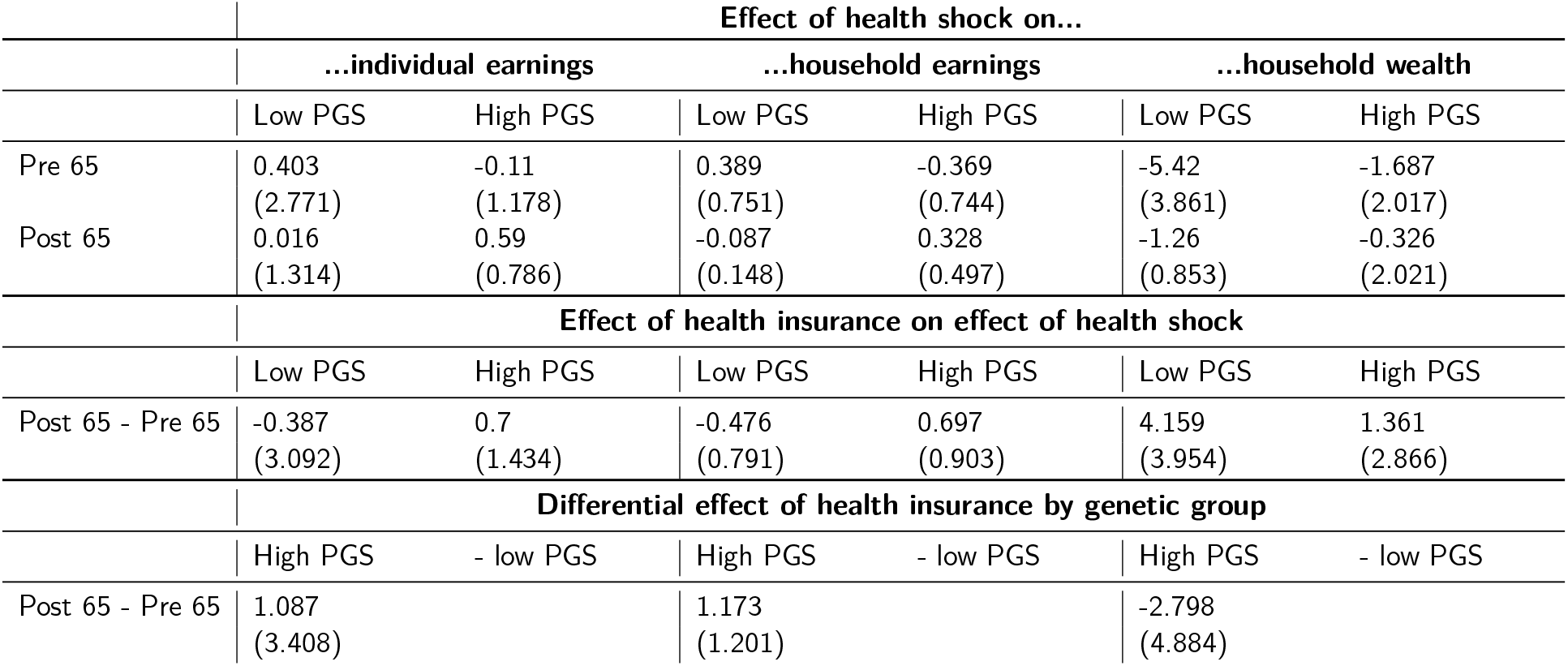
Summary of Placebo Checks for the Pre-65 Uninsured Subgroup for Income and Wealth

|  | Effect of health shock on... |  |  |  |  |  |
| --- | --- | --- | --- | --- | --- | --- |
|  | ...individual earnings |  | ...household earnings |  | ...household wealth |  |
|  | Low PGS | High PGS | Low PGS | High PGS | Low PGS | High PGS |
| Pre 65 | 0.403<br>(2.771) | -0.11<br>(1.178) | 0.389<br>(0.751) | -0.369<br>(0.744) | -5.42<br>(3.861) | -1.687<br>(2.017) |
| Post 65 | 0.016<br>(1.314) | 0.59<br>(0.786) | -0.087<br>(0.148) | 0.328<br>(0.497) | -1.26<br>(0.853) | -0.326<br>(2.021) |
|  | Effect of health insurance on effect of health shock |  |  |  |  |  |
|  | Low PGS | High PGS | Low PGS | High PGS | Low PGS | High PGS |
| Post 65 - Pre 65 | -0.387<br>(3.092) | 0.7<br>(1.434) | -0.476<br>(0.791) | 0.697<br>(0.903) | 4.159<br>(3.954) | 1.361<br>(2.866) |
|  | Differential effect of health insurance by genetic group |  |  |  |  |  |
|  | High PGS | - low PGS | High PGS | - low PGS | High PGS | - low PGS |
| Post 65 - Pre 65 | 1.087<br>(3.408) |  | 1.173<br>(1.201) |  | -2.798<br>(4.884) |  |

**Table D.16:** Summary of Placebo Checks for the Pre-65 Uninsured Subgroup for Retirement, Relationship Status and Out-of-Pocket Medical Expenditures

|  | Effect of health shock on... |  |  |  |  |  |
| --- | --- | --- | --- | --- | --- | --- |
|  | ...retirement |  | ...relationship status |  | ...OOP medical expenditures |  |
|  | Low PGS | High PGS | Low PGS | High PGS | Low PGS | High PGS |
| Pre 65 | 0.212<br>(0.265) | 0.094<br>(0.093) | 0.081<br>(0.073) | 0.044<br>(0.063) | 2.384***<br>(0.736) | 1.607***<br>(0.571) |
| Post 65 | 0.015<br>(0.093) | 0.022<br>(0.13) | 0.01<br>(0.01) | 0.041<br>(0.04) | 0.26<br>(0.658) | -1.27<br>(0.873) |
|  | Effect of health insurance on effect of health shock |  |  |  |  |  |
|  | Low PGS | High PGS | Low PGS | High PGS | Low PGS | High PGS |
| Post 65 - Pre 65 | -0.197<br>(0.28) | -0.071<br>(0.16) | -0.071<br>(0.07) | -0.003<br>(0.074) | -2.123**<br>(1.008) | -2.878***<br>(1.056) |
|  | Differential effect of health insurance by genetic group |  |  |  |  |  |
|  | High PGS | - low PGS | High PGS | - low PGS | High PGS | - low PGS |
| Post 65 - Pre 65 | 0.125<br>(0.322) |  | 0.068<br>(0.102) |  | -0.754<br>(1.457) |  |

**Table D.17:** Summary of Placebo Checks for the Pre-65 Uninsured Subgroup for Mortality within 2 and 5 years

|  | **Effect of health shock on mortality within...** |  |  |  |
| --- | --- | --- | --- | --- |
|  | **...2 years** |  | **...5 years** |  |
|  | Low PGS | High PGS | Low PGS | High PGS |
| Pre 65 | 0.004<br>(0.015) | 0.015<br>(0.09) | -0.046<br>(0.061) | -0.113<br>(0.119) |
| Post 65 | -0.001<br>(0.011) | -0.074<br>(0.065) | 0.204<br>(0.179) | 0.009<br>(0.103) |
|  | **Effect of health insurance on effect of health shock** |  |  |  |
|  | Low PGS | High PGS | Low PGS | High PGS |
| Post 65 - Pre 65 | -0.005<br>(0.025) | -0.089<br>(0.113) | 0.25<br>(0.191) | 0.122<br>(0.154) |
|  | **Differential effect of health insurance by genetic group** |  |  |  |
|  | High PGS | - low PGS | High PGS | - low PGS |
| Post 65 - Pre 65 | -0.085<br>(0.116) |  | -0.129<br>(0.245) |  |

Following these insights, we model the demand side of a health insurance market with agents who are heterogeneous in two dimensions: exogenous health risk and a moral hazard parameter governing the behavioral response towards being insured.

## E.1 Setup

Consider two time periods: in period 1, all agents are healthy^18^ and must make two decisions: whether to be insured, and how much effort to invest in reducing the probability of a health shock. More effort leads to a lower probability of falling sick, given an individual baseline level of risk. In period 2 the health risk realizes, with a probability mediated by the above-mentioned health-enhancing effort, and individuals can either be sick (*S*) or healthy (*H*). In case of sickness, insured agents receive the agreed-upon coverage 1 − *τ* and pay the premium *p*.^19^ Uninsured agents pay the full medical treatment. In the healthy state of the world, insured agents still pay the premium while uninsured keep their full income for consumption.

Utility depends on consumption *c* and the health state:

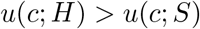

Utility is increasing and concave in consumption, i.e. *u′*(*c*;) > 0, *u″*(*c*; ·) ≤ 0. Utility from no consumption at all is 0 irrespective of the state of health, *u*(0; ·) = 0 and utility in consumption satisfies the Inada conditions^20^. The agent’s consumption is equal to the money available for consumption goods, i.e. a fixed income *y* which he receives in every period, less potentially insurance premia *p* and medical expenses *m* in case of sickness. Exerting health-enhancing effort *μ* is reducing his utility in the period it is exerted, which we phrase as health-enhancing effort cost. We assume effort costs to be additively separable from consumption utility and to be increasing and convex in the effort, i.e. *e′*(*μ*; ·) > 0, *e″*(*μ*; ·) > 0, *e*(0, ·) = 0. The effort costs are governed by the agent’s moral hazard type *g*. This type can be interpreted as some characteristics of the individual, which make it harder for him to refrain from harmful behavior such as smoking and eating unhealthy food or similarly to engage in health-enhancing behavior such as exercising. In the current setting, we consider the moral hazard type *g* to be proxied by an agent’s genotype, but it could also be interpreted as self-discipline or another individual characteristic. Higher predisposition to unhealthy behavior *g* leads to higher effort costs given the same level of effort, ∀*μ*: *e*(*μ*; *g′*) > *e*(*μ, g*) if *g′* > *g*. Regardless of whether the agent knows his genetic make-up, we assume that he is at least partly aware of his general type in the sense that he knows how easy it is for him to exert self-discipline.

The benefit of health-enhancing effort is the reduced probability of getting sick in the second period, which depends linearly on effort *λ* = *λ*_0_ − *μ*. The ex-ante risk type of the agent *λ*_0_ is equivalent to this probability, if no effort is exerted. The linear specification naturally bounds the maximal effort to *μ* ∊ [0, *λ*_0_], which raises the question of boundary solutions, which we will address later when going through the steps of the model.^21^

## E.2 The optimal health-enhancing effort

Since the game is finite, we solve it by backward induction. There is no decision to be made in the second period, the uncertainty about the health state just unfolds and the agent consumes what is left from his budget after paying his potential bills on insurance and medical expenditures.^22^ In the first period, the agent decides about whether to be insured *I* or not, as well as the health-enhancing effort *μ* he is willing to exert. Hence, he maximizes the present value of his utility stream:

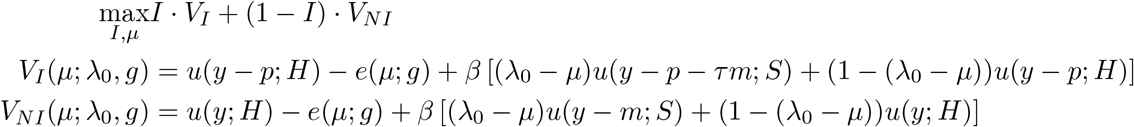

where *V_I_* is the value function when insured and *V_NI_* the value function when not insured, *β* is the time discount factor, *y* is income, *p* is the insurance premium, *m* is the medical expenditure, *τ* is the coverage rate, *H* and *S* are the healthy and sick state respectively, *μ* is the amount of health-enhancing effort, *g* is the (genetic) moral hazard type, and *λ*_0_ is the health risk type.

### 1 Proposition 1

*The agent exerts at least as much health-enhancing effort μ if he is insured compared to if he is not insured.*

Proof: Assuming an interior solution, if insured, the agent chooses optimal health-enhancing effort 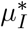 according to:

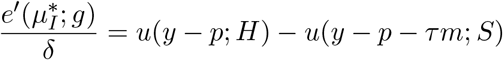

If uninsured, the FOC can be rearranged to:

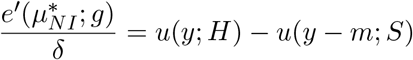

Now irrespective of the specific form of the utility function,^23^ 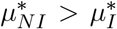. To see this more clearly consider subtracting (4) from (5):

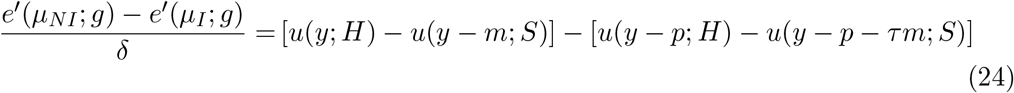

Effort in the uninsured state is larger than in the insured state, if the RHS of this expression is larger than zero (since effort costs are increasing and convex), which can be rearranged to:

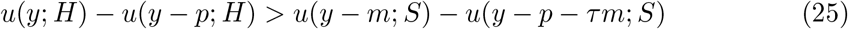

The LHS of this inequality is always positive, since *y* > *y* − *p*, while the RHS must be negative, since *m* > *p* + *τm*, otherwise, no agent ever chooses the insurance.^24^ Consequently, the above inequality will always hold in an interior optimum.

Now, let’s consider the boundary solutions. Since the marginal gains of exerting one extra unit of effort are constant, this consideration is rather simple. Suppose marginal cost at zero effort is higher than marginal gain in the case of no insurance: 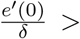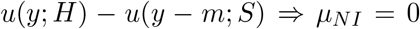.Then, we know that it must also be larger than marginal gain in case of insurance, i.e. *μ_I_* = 0. In this case, the health-enhancing effort is the same in both cases and the agent does not gain anything from being given the possibility to exert it. If the marginal cost at zero effort is below marginal gain in case of no insurance, but above marginal gain in case of insurance, the effort will be positive if not insured and thus higher than if not insured.

On the other bound, if marginal cost at exerting *λ*_0_ effort is lower than marginal gain if insured, then the agent will exert maximal effort in both cases. Finally, the effort will again be higher in case of no insurance compared to insurance, if marginal costs *e′*(*λ*_0_) is lower than marginal gain in case of no insurance, but higher in case of insurance, *μ_I_ < μ_NI_* = *λ*_0_. □

For the remaining part, we will assume an interior solution.

### Proposition 2

*Higher moral hazard parameter g leads to*

- *a smaller difference in health-enhancing effort μ between an agent having insurance and an agent not having insurance, all else equal.*
- *lower levels of effort when being insured and when not being insured respectively, compared to an agent with lower g and all else equal.*

Proof: As already mentioned, we assume that ∀*μ*: *e*(*μ*; *g′*) > *e*(*μ, g*) if *g′* > *g* and ∀*g*: *e*(0, *g*) = 0. Therefore it must be true that effort costs are supermodular in effort and genetic predisposition, i.e. 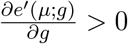 Consider (4) and (5) for two different values of genetic predisposition *g′* > *g*. Since the RHS of both equations is independent of *g*, it follows from the assumptions that *μ_I_*(*g*) > *μ_I_*(*g′*) and *μ_NI_*(*g*) > *μ_NI_*(*g′*). Moreover, note that the RHS of equation (24) is independent of the moral hazard parameter g. Hence, it must hold in equilibrium that:

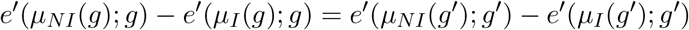

But since *μ_NI_*(*g*) > *μ_I_*(*g*), we know that *e′*(*μ_NI_*(*g*); *g*) − *e′*(*μ_I_*(*g*); *g*) < *e′*(*μ_NI_*(*g′*); *g′*) − *e′*(*μ_I_*(*g′*); *g′*). Consequently, it must hold that *μ_NI_*(*g*) − *μ_I_*(*g*) > *μ_NI_*(*g′*) − *μ_I_*(*g′*). □

Intuitively, Proposition 2 means, that agents for whom it is harder to engage in healthy or disengage in unhealthy behavior, e.g. starting to exercise or quit smoking, will react less to an increasing benefit to do so. Thus, a higher moral hazard parameter *g* coincides with less moral hazard because there is less leeway for the individual to adjust effort to insurance coverage. A higher moral hazard parameter also implies a lower effort of the agent, in both states of the world. This agent is *more* risky from the perspective of the insurance *ex-ante as well as ex-post* despite showing less reaction to insurance coverage.

Our empirical results outlined above can be considered an empirical counterpart to this proposition, leveraging the occurrence of a health shock to make the situation more salient to the individual.

### Proposition 3

*Given genetic predisposition g, an agent with higher risk type λ*_0_ *has a higher net benefit from being insured*.

Proof: Let us denote the present values of the optimal health-enhancing effort level given the moral hazard type *g* as 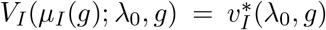, 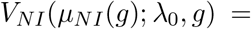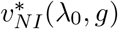. In period 1, the agent will choose to be insured, if 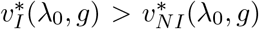 Both expressions consist of parts that are dependent on the optimal health preventing effort and those that are not:

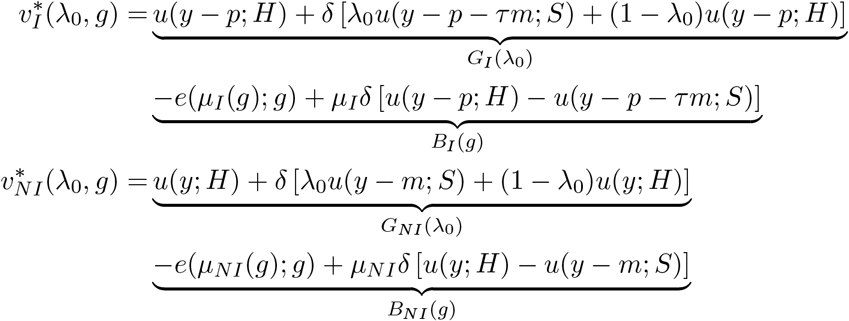

From equation (25), it follows that 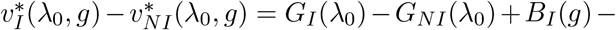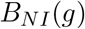 is linearly increasing in health risk *λ*_0_. □

### Proposition 4

*Higher genetic predisposition g leads to higher adverse selection on risk type*.

Proof: For the lowest risk type *λ*_0_ = 0 there is no effort he can exert to improve his health prospect, and he will never choose to be insured:

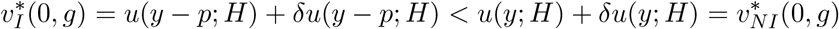

Note that a risk type 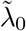, who is indifferent between insurance and no insurance with zero health-enhancing effort, is *not* choosing the insurance with optimal effort: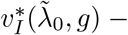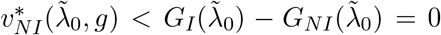 Hence there exists a new threshold 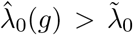 above which all agents choose to be insured given genetic predisposition.

From Proposition 4, we know that a higher moral hazard parameter *g* leads to a higher net benefit in being insured. To evaluate how the threshold risk type 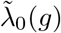 is changing in the moral hazard parameter, we can use the implicit function theorem on equation (8):

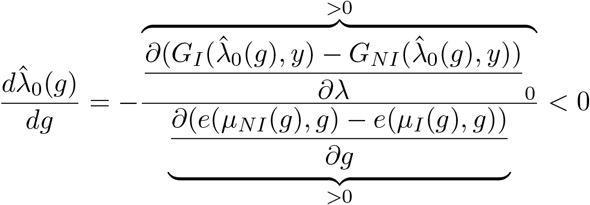

Consequently, if we have two groups of agents with genotype *g′* > *g*, then adverse selection will be less severe for the group with the higher moral hazard parameter 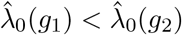 and the lower reaction of effort to insurance. □

Note that because we did not further specify the insurance contract offered by the insurer, nothing prevents some thresholds to be above 1, i.e. no agent of a given genetic predisposition might choose the insurance. We assume that without health-enhancing effort, the highest risk type *λ*_0_ = 1 would always like to choose the insurance, i.e.^25^:

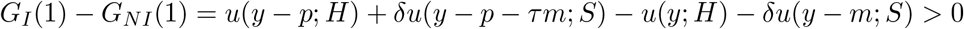

This is equivalent to stating that there exists a 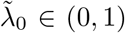. Now allowing for health-enhancing effort, it will depend on *g*, whether insurance is preferred to no insurance:

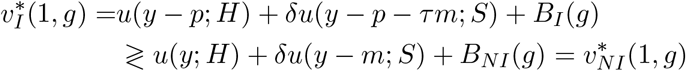

Finally, if 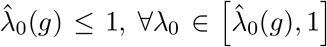 the agent is choosing the insurance and 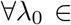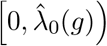 the agent is not choosing the insurance, all else equal. This is the usual adverse selection result: Agents with a higher risk of falling sick inflict more costs to the insurance, but also have a higher valuation for insurance. If 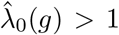 which might happen for a very low level of *g*, we have the extreme case that no agent of a given moral hazard parameter *g* is choosing to be insured.

The following picture summarizes the relationship:

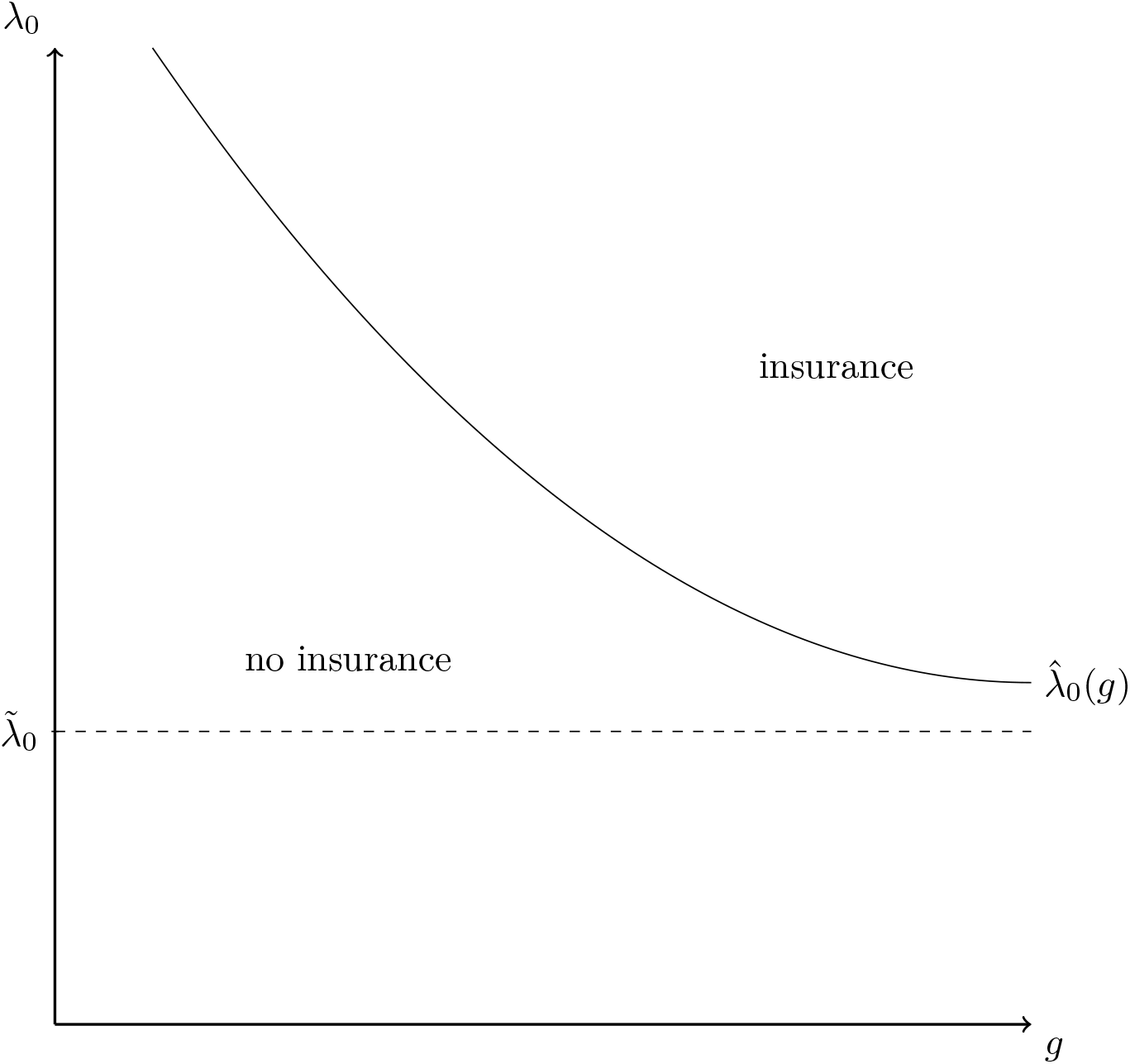

Over a population with independently distributed risk and moral hazard types we should expect to see agents of low risk type 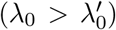 choosing an insurance only if they have high costs of self-discipline (*g > g′*). Agents with high exogenous health risks should choose an insurance irrespective of their moral hazard type. Consequently, within a self-selected population with insurance we should observe a positive correlation of risk type and degree of moral hazard (or a negative correlation of risk type and moral hazard parameter g), despite advantageous selection on moral hazard, i.e. despite the fact that within each risk group, agents with the largest moral hazard potential do not choose the insurance.

While exogenous risk type *λ*_0_ is an interesting variable to look at theoretically, it is neither the realized risk of the insured or uninsured agent. To compare the realized risk with and without health-enhancing effort, it is useful to divide the indirect utility when insured and uninsured into its components. We define 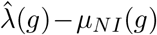 as the threshold exante risk of an insured agent, which is what researchers usually consider the exogenous risk type on which adverse selection occurs.

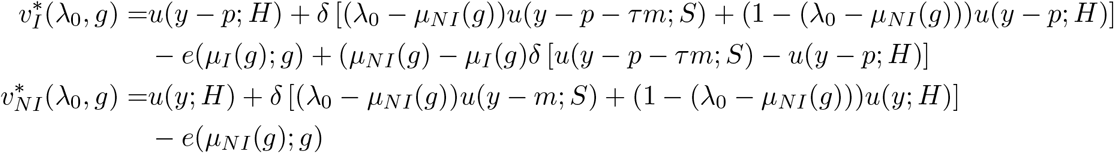

From this reformulation, it is easy to see that the relatively higher attractiveness of no insurance does not only result from the lower realized health risk *λ*_0_ − *μ_NI_*(*g*) compared to exogenous risk type *λ*_0_. Also, the behavioral reaction to being covered^26^ makes insurance *less* attractive. Moral hazard drives a wedge between the health risk if an agent is covered and the health risk if he is not covered, which can be interpreted as the agent’s differential attempt to prevent the bad state of the world himself. Consequently, it must be true that 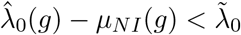, but 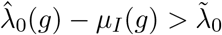, so health-enhancing effort leads to an increase in realized risk among the insured agents given the same contract. This, in general, makes the adverse selection problem more severe. An insurer offering a fixed contract thus should increase premia in a world with a behavioral response of the agent compared to one without a behavioral response. which would exclude more agents from being insured. Notice, however, that not being insured has the positive side effect of the agents partly reduce risk themselves and staying healthy more likely.

## Footnotes

1 Response rates vary between 80% and 90% (HRS, 2017*a*). Every 6 years, a new 6-year birth cohort of participants is enrolled (HRS, 2017*b*). It is sponsored by the National Institute on Aging (grant number NIA U01AG009740) and is conducted by the University of Michigan (HRS, n.d.). The first core interview with every participant is conducted face-to-face, and all follow-up interviews are either face-to-face or over the phone (HRS, 2017*b*).

2 See Section C.2 in the Appendix for more information on the construction of the study sample.

3 Martin et al. (2017) convincingly show that past attempts to use polygenic scores to compare racial or ethnicgroup achievements were scientifically flawed. Sadly, this reduces the external validity of any study that uses polygenic scores informed by GWAS of white participants. It might also exacerbate existing health disparities across ethnicity and hamper the potential for scientific knowledge and innovation to improve everyone’s lives (Martin et al., 2019). Thankfully, multi-ancestry GWAS are becoming more common, but not yet for smoking behavior (Peterson et al., 2019).

4 The results do not change if these individuals are included.

5 As a robustness check, shown in Appendix Section D.3.3, we also use the polygenic score provided by the HRS (Ware, Schmitz and Faul, 2017) for the smoking phenotype “regular smoking” (having smoked more than 100 cigarettes throughout one’s life). This score is constructed as a weighted sum of the genotype over the 779,538 SNPs that overlap between the HRS genetic database and a 2010 GWAS meta-analysis conducted by the Tobacco and Genetics Consortium (The Tobacco and Genetics Consortium et al., 2010). See Section C.3.5 in the Appendix for more information on the construction of the PGS.

6 We initially ran the analysis separately for each tertile of the PGS distribution, as shown in Appendix Section D.3.1, but the results are concentrated in the lower part of the distribution of the PGS, and therefore for simplicity we consider only high and low PGS. Results are robust to splitting the same above and below the median PGS score, as shown in Appendix Section D.3.2.

7 This slight gender imbalance is a feature of our sampling strategy and not of the PGS, which is balanced across genders in the original paper (Liu et al., 2019). None of the results are driven by gender differences, which are always controlled for in the analysis. Appendix Section D.1 provides additional descriptive information about the study sample that experienced a health shock.

8 See Appendix Section C.5 for a derivation of these effects.

9 Notice that the magnitude of this coefficient (0.09) is similar to the ones estimated for high polygenic score individuals (−0.108 and −0.13), but it is much more precisely estimated. The main reason behind smaller standard errors for the estimated effect of health shock after 65 for the low polygenic score individuals is the negative covariance 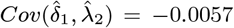. See Appendix Table 7 for the variance-covariance matrix of the estimated coefficients of equation 2. *λ*2 captures the effect of the health shock in the uninsured population, and *δ*1 the incremental effect of that same health shock in the uninsured population after the age of 65.

10 See for instance Anastasi (1958); Haldane (1946); Ottman and Rao (1990); Plomin (1990); Plomin, DeFries and Loehlin (1977).

11 Calculated by HRS as the sum of a word recall task (Total Recall Index) and a counting, naming, and vocabulary task (Mental Status Index).

12 Notice that some of these variables are not quite predetermined, baseline characteristics of the individuals. Some might actually be mediators of the overall effect, or “bad controls” in the terminology of Angrist and Pischke (2008), especially income. We still report the results for completeness, but caution the reader prone to causal interpretations. In this regard, genetic variants represent the ideal measure of heterogeneity of treatment effects: they are fixed since conception, immutable, identically measured across the whole human species, and indexing plausible biological channels which are increasingly studied and documented by a wide range of disciplines. In Appendix D.3.4, we report the results of using different polygenic scores as a proxy for *g_i_*. Moral hazard heterogeneity is also detected in the group having a high PGS for educational attainment or cognitive abilities (Lee et al., 2018), and a low PGS for risk tolerance (Karlsson Linnér et al., 2019) and a low PGS for non-cognitive skills (Demange et al., 2020). On the other side, there is no evidence of heterogeneity of moral hazard by the PGS for Body-Mass-Index (Yengo et al., 2018) or the PGS for cigarettes per day (Liu et al., 2019). These results suggest that potential mechanisms for the observed heterogeneity should include not only biological channels related to nicotine addiction, but also cognitive processes related to risky and strategic behavior, which are closely related to the concept of moral hazard.

13 See number of observations in Appendix Table 5.

14 For example, see the model in Appendix E.

15 A base *pair* is set of two bases, with A always pairing with T, and C always pairing with G.

16 This model was initially drafted by Regina Seibel, who was working on the project as a research assistant. An abridged version of the model is reported here with her permission.

17 This change in risky health behaviors due to the anticipation of being able to afford health care in the future has been dubbed *ex-ante* moral hazard. *Ex-post* moral hazard refers to an increase in the use of medical care—such as doctor visits or medicines—following health insurance coverage.

18 In other words, we do not consider the case of pre-existing conditions.

19 To start off with a model as simple as possible, we abstract from the supply side and just assume that the insurer offers a fixed contract with premium *p* and coverage rate 1 *τ*. A special case would be complete insurance with *τ* = 0.

20 Inada conditions: lim_*c* → 0_ *u′*(*c*; •) = ∞, lim_*c* → ∞_ *u′*(*c*; •) = 0

21 Another way to model the problem would be to allow for a more general functional form *λ*(*μ*). It is reasonable to assume risk to be decreasing in effort, i.e. *λ′*(*μ*). However, assumptions about convexity or concavity of the function are less straighforward and would require further supporting evidence. One could find examples supporting both increasing and decreasing marginal effects of effort on risk. Note that the results are robust to slight convexity or concavity. Extreme concavity might rule out an interior solution - the problem becomes a choice between no effort and maximal effort - while extreme convexity renders Proposition 1 is not as clear-cut.

22 One could think additionally modeling the decision about the size of the medical expenditures, for example a choice between an expensive or a cheap treatment. This is what Einav et al. (2013) consider as moral hazard. Insured agents are more likely to choose the expensive treatment.

23 In particular, irrespective of the interaction between health and income in determining the agent’s utility, i.e. utility being sub-modular or supermodular in health and consumption.

24 If *m* ≤ *p* + *τm*, the insurance renders the agent weakly worse off in the sick state, while making him strictly worse off in the healthy state, since he has to pay the premium. A rational agent, irrespective of his idiosyncratic health risk, would never choose such an insurance.

25 This is an assumption that needs to be justified by looking at the firm side later and solve for the market equilibrium of the problem

26 Note that *μNI δ*(*u*(*y − p*; *H*) − *u*(*y − p τm*; *S*)) − *e*(*μNI* (*g*); *g*) < *μ_I_ δ*(*u*(*y − p*; *H*) − *u*(*y − p − τm*; *S*)) − *e*(*μ_I_* (*g*); *g*), since *μ_I_* (*g*) is the maximum of the problem of the insured agent.

